# ASO-enhancement of *TARDBP* exitron splicing mitigates TDP-43 proteinopathies

**DOI:** 10.1101/2024.07.22.604579

**Authors:** Takuma Yamagishi, Shingo Koide, Genri Toyama, Aya Washida, Yumi Yamada, Ryutaro Hanyu, Ekaterina Nadbitova, Yuka Mitsuhashi Koike, Takuya Konno, Tomohiko Ishihara, Taisuke Kato, Osamu Onodera, Akihiro Sugai

## Abstract

Amyotrophic lateral sclerosis and frontotemporal lobar degeneration are fatal neurodegenerative diseases characterized by pathological aggregation and nuclear functional loss of TDP-43^1,2^. Current therapies inadequately address this core pathology^3,4^, necessitating innovative approaches that target aggregation while preserving TDP-43’s essential functions. Here we demonstrate that enhancing the splicing of the *TARDBP* exitron—a cryptic intron encoding the aggregation-prone intrinsically disordered region (IDR) of TDP-43^5,6^— effectively mitigates TDP-43 pathology. This exitron splicing event, directly regulated by nuclear TDP-43^7–9^, suppresses the expression of IDR-containing TDP-43 isoforms and generates IDR-spliced-out TDP-43 isoforms^7,9,10^ (which we term “IDRsTDP”). Our findings reveal that IDRsTDP, known to heterodimerize with full-length TDP-43^10^, inhibits TDP-43 aggregation by suppressing IDR-mediated clustering and enhances TDP-43 clearance via chaperone-mediated autophagy. In disease states, however, impaired nuclear TDP-43 function disrupts exitron splicing, leading to increased levels of IDR-containing TDP-43^9,11^ and reduced levels of IDRsTDP, exacerbating aggregation and nuclear dysfunction^6,12–17^. By identifying HNRNPA1 and HNRNPC as key repressors of *TARDBP* exitron splicing, we designed antisense oligonucleotides (ASOs) to block their binding and restore splicing. These ASOs suppressed TDP-43 pathology and neurodegeneration in both neuronal cell models with impaired nuclear transport and a mouse model of proteasome dysfunction-induced TDP-43 proteinopathy. Our strategy, by rescuing the impaired autoregulatory pathway, inhibits the pathological cycle of TDP-43 aggregation and nuclear dysfunction, offering a promising avenue for treating these currently intractable neurodegenerative diseases.

## Main

Sporadic amyotrophic lateral sclerosis (ALS) and frontotemporal lobar degeneration (FTLD) are devastating neurodegenerative diseases characterized by pathological aggregation and mislocalization of TDP-43, a nuclear DNA/RNA-binding protein essential for post-transcriptional regulation^1,2,7,18,19^. Current therapeutic strategies for ALS and FTLD focus on symptom management or addressing downstream consequences of TDP-43 pathology, including excitotoxicity or inflammation^3,4^. However, these therapies do not directly target the core pathogenic mechanisms driving TDP-43 aggregation and dysfunction.

Our study focuses on a promising therapeutic target: the intrinsic autoregulatory mechanism controlling TDP-43 expression. TDP-43 regulates its own levels through a negative feedback loop involving the alternative splicing of its pre-mRNA transcript^7,8^. Specifically, nuclear TDP-43 promotes the splicing of an exitron—a cryptic intron located within exon 6 of *TARDBP*—which encompasses the entire coding region for its intrinsically disordered region (IDR) (Fig. 1a)^7,9,11^. As the IDR is critical for TDP-43’s propensity to aggregate^5,6^, exitron splicing effectively removes this aggregation-prone region from over half of the transcripts, highlighting a sophisticated mechanism for maintaining TDP-43 homeostasis^6,11,14^. In TDP-43 proteinopathies, however, the autoregulatory mechanism is impaired by nuclear TDP-43 dysfunction^6,9,13^ and ALS-causing TDP-43 mutations^20–22^. Consequently, this leads to an overabundance of IDR-containing transcripts and increased TDP-43 aggregation^11,14^.

**Fig. 1.**
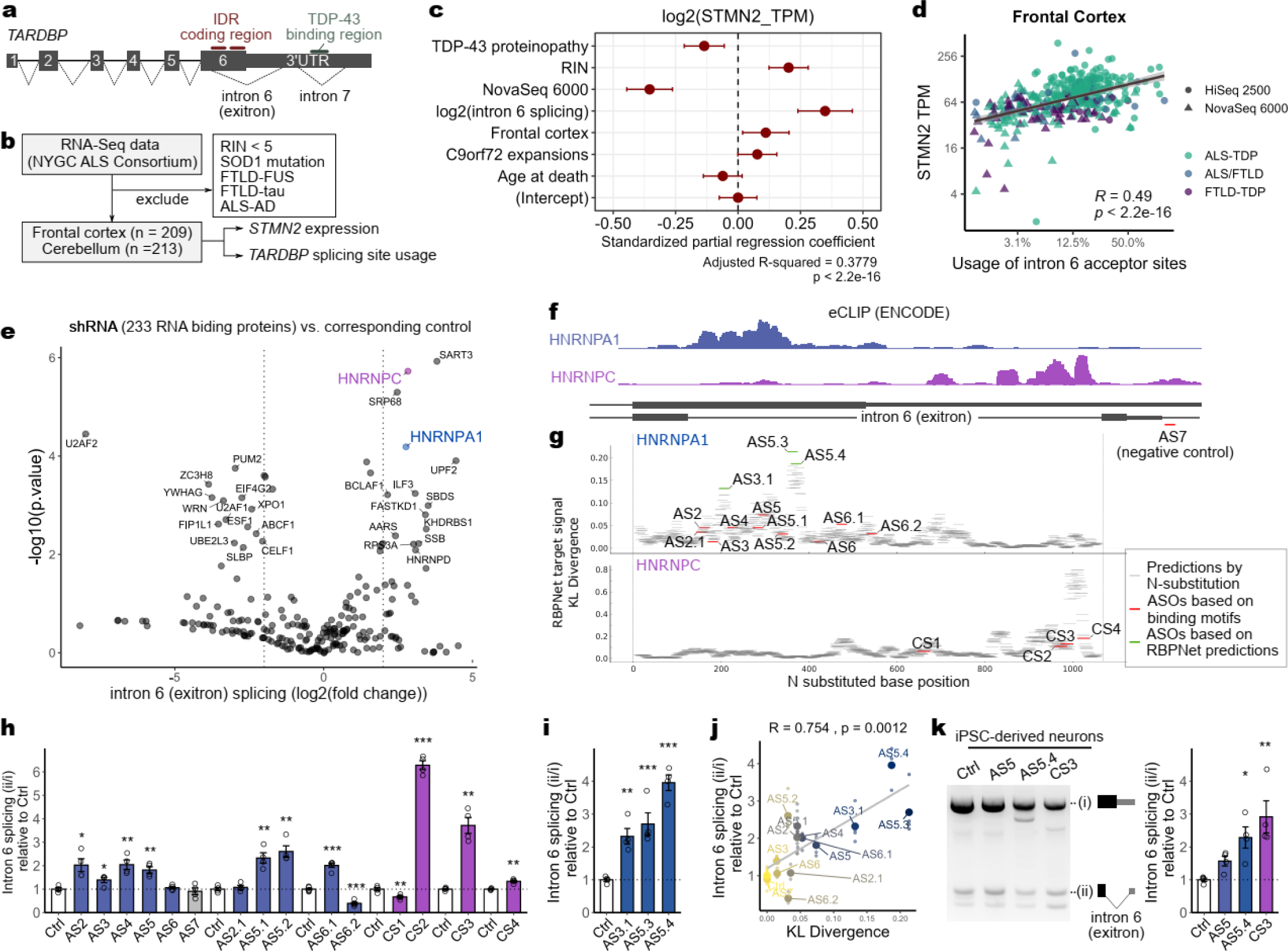
*TARDBP* exitron splicing is reduced by TDP-43 dysfunction and enhanced by ASO-mediated HNRNPA1/C binding inhibition. **a,** Schematic of *TARDBP* pre-mRNA showing the relationship between exitron and IDR coding regions. **b-d,** Analysis of NYGC ALS Consortium RNA-seq data. **b,** Analysis criteria. **c,** Multiple regression analysis with log-transformed *STMN2* expression as the dependent variable. **d,** Correlation between exitron splicing and *STMN2* expression in the frontal cortex of patients with TDP-43 proteinopathies. **e,** RNA-binding proteins (RBPs) altering exitron splicing, based on ENCODE RNA sequencing data from HepG2 cells with knockdown of 233 RBPs. **f,** ENCODE eCLIP data for HNRNPA1 (ENCFF706CRY) and HNRNPC (ENCFF774JCA). **g,** RBPNet simulation of ASO-mediated binding inhibition of HNRNPA1 and HNRNPC. Larger KL divergence indicates greater inhibition by the ASO sequence. **h,i,** Intron 6 (exitron) splicing analysis after cycloheximide (CHX) treatment in HEK293T cells transfected with ASOs targeting HNRNPA1/C binding regions. **j,** Correlation between predicted HNRNPA1 binding inhibition efficiency (KL divergence from RBPNet) and ASO-mediated exitron splicing promotion. **k,** RT-PCR showing ASO effects on exitron splicing in human iPSC-derived neurons. Data are presented as means ± SEM. ***p < 0.001, **p < 0.01, *p < 0.05.

To further establish the clinical relevance of *TARDBP* exitron splicing, we analyzed RNA sequencing data from a cohort of healthy controls and patients with TDP-43 proteinopathies (GSE153960; Fig. 1b). Notably, we observed a strong positive correlation between *TARDBP* exitron splicing rates and *STMN2* expression, a sensitive marker of nuclear TDP-43 functions^23–25^ (β = 0.35, p = 8.55e-10; Fig. 1c). This correlation was particularly prominent in the frontal cortex, a brain region highly susceptible to TDP-43 pathology (Fig. 1d, Extended Data Fig. 1a, b). Furthermore, exitron splicing rates in the healthy frontal cortex exhibited an age-dependent decline (Extended Data Fig. 1c), consistent with age being a primary risk factor for ALS and FTLD^26^.

Based on this compelling observation, we hypothesized that enhancing *TARDBP* exitron splicing could represent a novel therapeutic strategy for TDP-43 proteinopathies. By modulating this intrinsic regulatory pathway, we aim to reduce the burden of aggregation-prone TDP-43 and potentially mitigate the pathogenic cascade underlying these devastating diseases.

### ASOs targeting HNRNPA1 and HNRNPC enhance exitron splicing

To identify potential therapeutic targets for enhancing *TARDBP* exitron splicing, we sought to identify key RNA-binding proteins (RBPs) regulating this process. Analysis of ENCODE data from knockdown experiments targeting 233 RBPs revealed that several significantly affected *TARDBP* exitron splicing (Fig. 1e). We focused on HNRNPA1 and HNRNPC, which exhibited pronounced binding motifs flanking the 5’ and 3’ ends of the exitron, respectively (Extended Data Fig. 1d), suggesting a direct regulatory role. ENCODE eCLIP data confirmed the binding of these proteins to the *TARDBP* exitron (Fig. 1f). To directly assess their function, we used a minigene system containing the *TARDBP* exitron in HEK293T cells. Consistent with the ENCODE data, knockdown of either HNRNPA1 or HNRNPC significantly enhanced *TARDBP* exitron splicing (Extended Data Fig. 1e-i), supporting their role as splicing repressors.

Next, we designed morpholino ASOs targeting the binding peaks of HNRNPA1 and HNRNPC within the *TARDBP* pre-mRNA (Fig. 1g). Gratifyingly, seven ASOs targeting HNRNPA1 binding regions and three targeting HNRNPC regions effectively promoted exitron splicing (Fig. 1h, Extended Data Fig. 2a, c, d). To further enhance ASO efficacy, we leveraged RBPNet^27^, a machine learning algorithm, to predict sequences with improved ability to disrupt RBP binding (Fig. 1g). This approach yielded ASOs with significantly greater splicing-promoting effects compared to those designed solely based on binding site location (Fig. 1i, j, Extended Data Fig. 2b, e, f). Importantly, these ASOs also successfully promoted exitron splicing in human iPSC-derived neurons (Fig. 1k), highlighting their therapeutic potential.

### ASO-mediated enhancement of impaired *TARDBP* exitron splicing mitigates TDP-43 pathology

To investigate the therapeutic potential of targeting *TARDBP* exitron splicing in a disease-relevant context, we employed cellular models of TDP-43 proteinopathy based on knockdown of CSE1L/CAS, a factor essential for TDP-43 nuclear import that is reduced in the frontal lobes of FTLD-TDP patients^28,29^. CSE1L/CAS knockdown in SH-SY5Y cells recapitulated key pathological hallmarks observed in patient brains, including reduced nuclear TDP-43 levels and function, as evidenced by decreased *TARDBP* exitron splicing and diminished *STMN2* expression (Fig. 2a, b). These findings were consistent with our observations in human autopsy brain samples (Fig. 1d). Additionally, this model exhibited robust accumulation of insoluble TDP-43 (Fig. 2c), providing a platform to examine the effects of enhancing exitron splicing on TDP-43 aggregation.

**Fig. 2.**
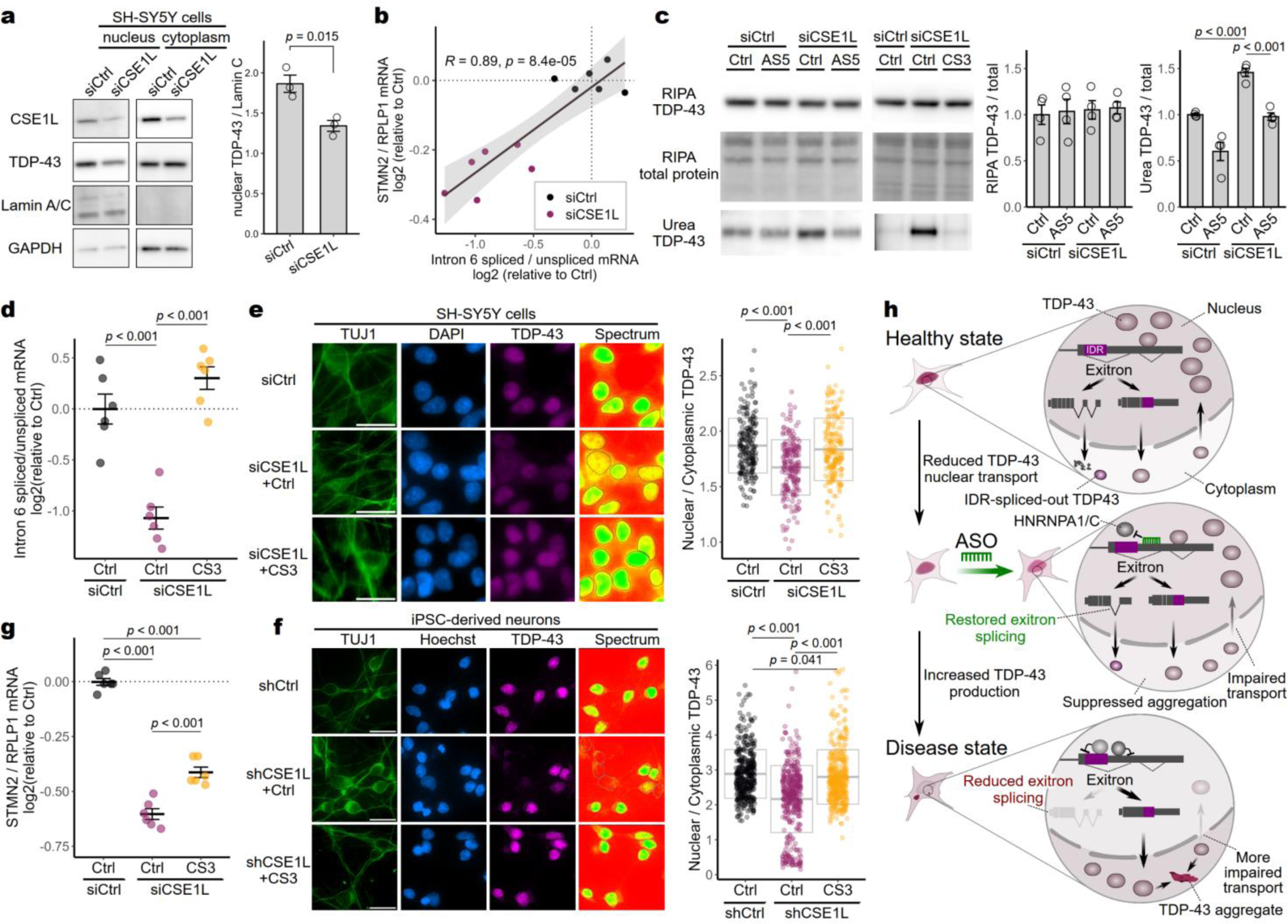
ASO-mediated restoration of *TARDBP* exitron splicing alleviates TDP-43 pathology induced by nuclear translocation defect. **a,** Western blot analysis of nuclear TDP-43 in SH-SY5Y cells following CSE1L knockdown. **b,** Correlation between *TARDBP* intron 6 (exitron) splicing and *STMN2* expression levels in SH-SY5Y cells treated with siCSE1L or control siRNA. **c,** Western blot analysis of TDP-43 in RIPA and Urea fractions of SH-SY5Y cells treated with siCSE1L and ASOs. **d,** Exitron splicing efficiency in SH-SY5Y cells treated with siCSE1L and ASOs. **e,f,** Immunofluorescence images showing TDP-43 localization in SH-SY5Y cells (**e**) and iPSC-derived neurons (**f**) following CSE1L knockdown and ASO treatment. Scale bar, 20 µm. g, *STMN2* expression levels of SH-SY5Y cells treated with siCSE1L and ASO, measured by qRT-PCR. **h,** Schematic illustrating the proposed mechanism by which restoration of exitron splicing mitigates TDP-43 proteinopathy. Data are presented as mean ± SEM (**ac,d,g)** or mean ± SD (**e,f**). Statistical analysis was performed using Student’s t-test (**a**), Pearson’s correlation test (**b**), ANOVA followed by Tukey’s multiple comparison test (**c-e**), and the Kruskal-Wallis test followed by Dunn’s multiple comparison test with Bonferroni correction (**f**).

We first confirmed that ASOs targeting either HNRNPA1 or HNRNPC binding regions effectively restored *TARDBP* exitron splicing in CSE1L/CAS-knockdown cells (Fig. 2d, Extended Data Fig. 3a-c). Remarkably, restoring exitron splicing in this model significantly mitigated the accumulation of insoluble TDP-43 without affecting soluble TDP-43 levels (Fig. 2c). Furthermore, directly inhibiting exitron splicing by targeting the splice junction was sufficient to induce insoluble TDP-43 accumulation in SH-SY5Y cells (Extended Data Fig. 3d), consistent with our previous findings in mouse spinal cord^11^. These results underscore the critical role of exitron splicing in preventing TDP-43 aggregation.

We next investigated whether restoring exitron splicing could rescue the nuclear TDP-43 depletion observed in our model. TDP-43 aggregation is thought to exacerbate its nuclear depletion through a vicious cycle, whereby aggregates sequester newly synthesized TDP-43 and interfere with nuclear import machinery^6,12,13,15–17^. This, in turn, may further reduce exitron splicing, leading to more aggregation and progressive loss of nuclear function^5,6,13,14^. Our previous observation that inhibiting exitron splicing in iPSC-derived neurons was sufficient to decrease nuclear TDP-43 supports this interconnected mechanism^11^.

Remarkably, ASO-mediated enhancing of exitron splicing significantly attenuated the reduction in the nuclear/cytoplasmic TDP-43 ratio induced by CSE1L/CAS knockdown in both SH-SY5Y cells (Fig. 2e, Extended Data Fig. 3e) and iPSC-derived neurons (Fig. 2f). Functionally, this restoration of nuclear TDP-43 by ASO treatment in CSE1L/CAS-knockdown SH-SY5Y cells partially rescued the expression of *STMN2* (Fig. 2g), a key downstream target of TDP-43^23–25^.

These results demonstrate that reactivating exitron splicing effectively breaks the pathological cycle of TDP-43 mislocalization and aggregation by mitigating TDP-43 aggregation and preserving nuclear TDP-43 function, even under conditions of impaired nuclear import (Fig. 2h).

### IDR-spliced-out TDP-43 isoforms suppress TDP-43 aggregation

To further understand the mechanism by which enhancing exitron splicing exerts its protective effects, we investigated the potential role of the alternative TDP-43 isoforms generated by this process^7,9,10^. Indeed, ASO (CS3) that promotes exitron splicing significantly increased the levels of IDR-spliced-out TDP-43 isoforms (hereafter referred to as “IDRsTDP”) (Extended Data Fig. 4a, b). The IDRsTDP lack the aggregation-prone IDRs but retain the N-terminal domain (NTD) (Fig. 3a). These isoforms can heterodimerize with full-length TDP-43 via the NTD^10,30–32^ (Extended Data Fig. 4c), leading us to hypothesize that the IDRsTDP might inhibit TDP-43 aggregation through liquid-liquid phase separation (LLPS), which is driven by increased local concentration of IDRs (Fig. 3a) ^33,34^.

**Fig. 3.**
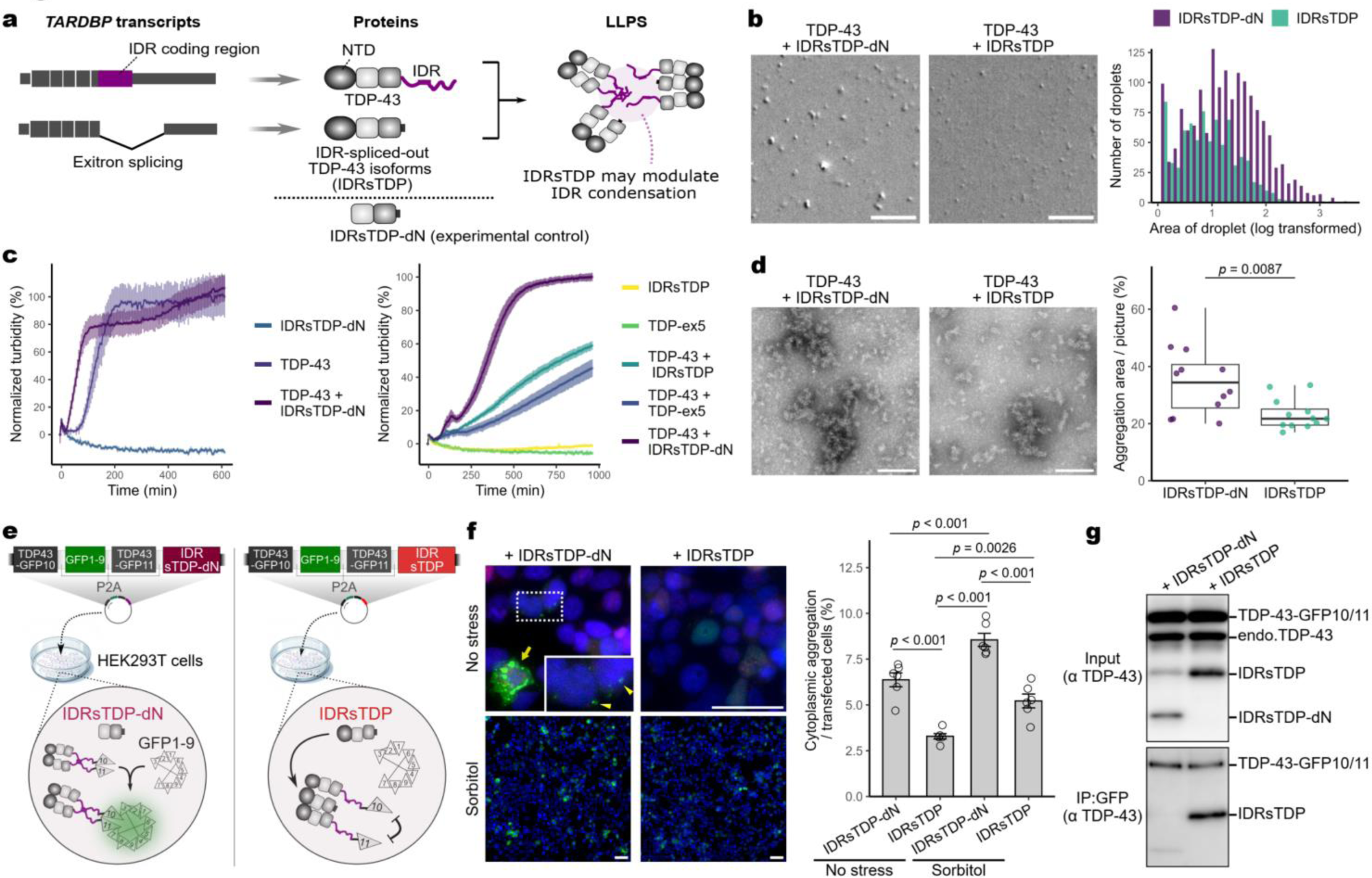
IDR-spliced-out TDP-43 isoforms (IDRsTDP) inhibits TDP-43 aggregation. **a,** Schematic diagram of the relationship between exitron splicing and IDR-spliced-out TDP-43 isoforms (IDRsTDP), and how IDRsTDP modulates IDR condensation. **b,** Droplet formation of TDP-43 with IDRsTDP or IDRsTDP-dN. **c,** Aggregation assays comparing TDP-43 alone and with IDRsTDP, IDRsTDP-dN, TDP-ex5. **d,** TEM images and quantification of aggregation area percentage. **e,** Schematic of the Split-GFP system experiment. **f,** Immunostaining of HEK293T cells transfected with Split-GFP constructs under normal and osmotic stress conditions (sorbitol). **g,** Immunoprecipitation with GFP-trap beads followed by Western blot analysis. Scale bars: 20 µm (**b**), 100 nm (**d**), 40 µm (**f**). Statistical analysis: Welch’s t-test (**d**), ANOVA with Tukey’s test (**f**).

To test this hypothesis, we purified full-length TDP-43, IDRsTDP, and a control IDRsTDP lacking NTD (IDRsTDP-dN) (Fig. 3a). In both in vitro LLPS and aggregation assays, IDRsTDP significantly reduced TDP-43 droplet formation and aggregation compared to IDRsTDP-dN (Fig. 3b-d). Notably, TDP-ex5, encoded up to exon 5 of *TARDBP* that also retains the NTD but lacks the IDR, similarly inhibited TDP-43 aggregation (Fig. 3c), further supporting the importance of the IDR lack and NTD in mediating this effect.

To validate these findings in a cellular context, we employed a Tripartite Split-GFP Complementation system in HEK293T cells, which allows for visualization of TDP-43 IDR proximity through fluorescence reconstitution^30,35^ (Fig. 3e). Cells expressing IDRsTDP displayed significantly fewer GFP granules compared to those expressing IDRsTDP-dN, indicating reduced TDP-43 IDR clustering (Fig. 3f). Furthermore, under osmotic stress conditions that typically induce TDP-43 mislocalization and aggregation^36^, IDRsTDP expression effectively suppressed cytoplasmic GFP aggregation. Co-immunoprecipitation experiments confirmed that this suppressive effect was mediated by direct interaction between IDRsTDP and TDP-43 through their respective NTDs (Fig. 3g). These results demonstrate that IDRsTDP can effectively inhibit TDP-43 aggregation in cell.

### IDRsTDP reduces full-length TDP-43 levels via chaperone-mediated autophagy

We next investigated the effects of IDRsTDP on TDP-43 protein levels, given the interaction between IDRsTDP and TDP-43 via their NTDs. Transient overexpression of IDRsTDP reduced endogenous full-length TDP-43 levels, suggesting accelerated degradation (Fig. 4a). This degradation mechanism appears distinct from the C-terminal fragment-mediated degradation observed in TDP-43 proteinopathies (Extended Data Fig. 5a)^37^, suggesting a novel regulatory pathway.

**Fig.4. IDRsTDP.**
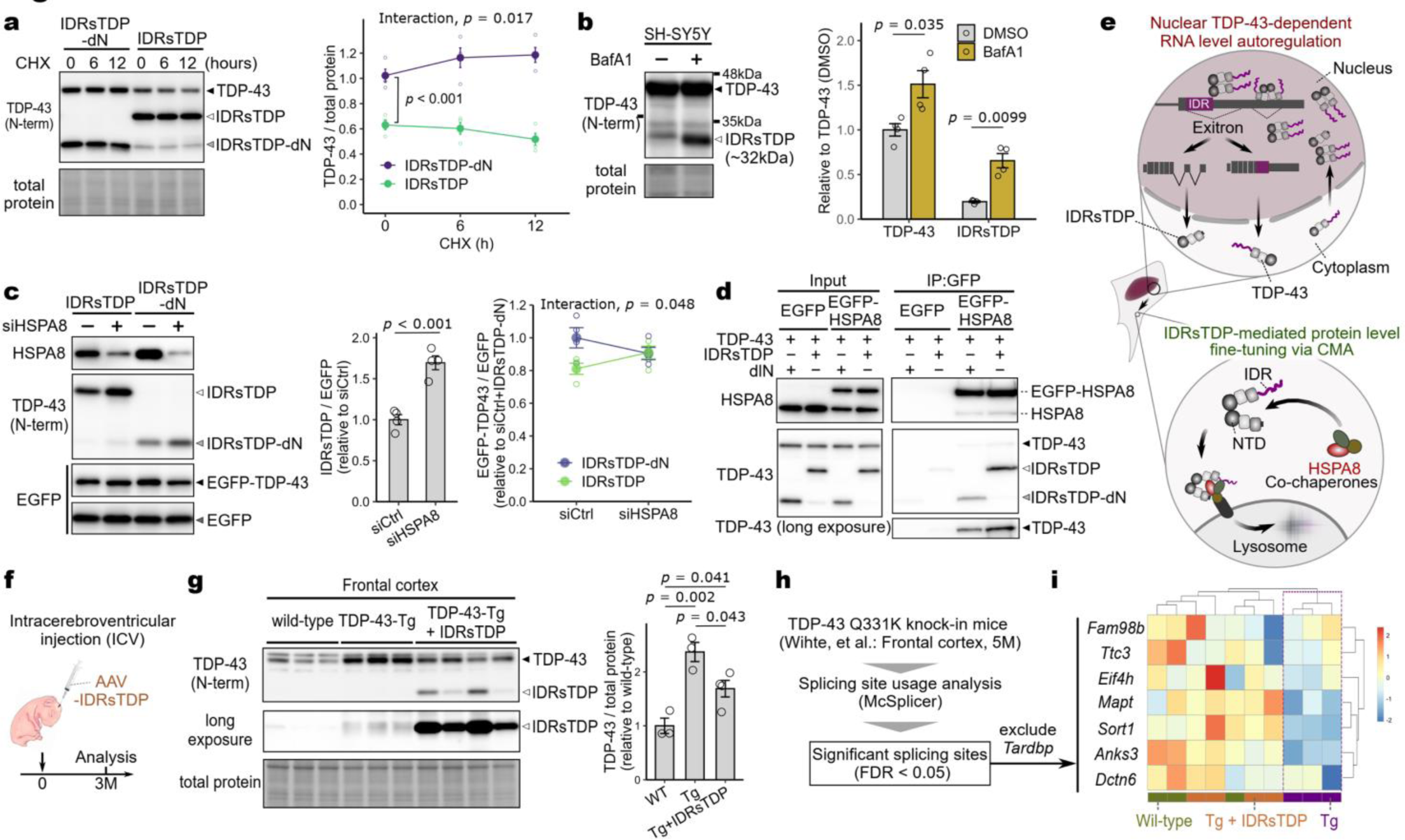
suppresses excess TDP-43 via chaperone-mediated autophagy (CMA) and abnormal RNA metabolism in TDP-43 Tg mice. **a,** Stability of endogenous TDP-43 in HEK293T cells expressing IDRsTDP or IDRsTDP-dN following CHX treatment. **b,** TDP-43 and IDRsTDP expression levels in SH-SY5Y cells treated with bafilomycin A1 (BafA1). **c,** Effect of knockdown of HSPA8 on IDRsTDP and EGFP-TDP-43 levels in HEK293T cells expressing IDRsTDP or IDRsTDP-dN. EGFP serves as a transfection control. **d,** Co-immunoprecipitation of EGFP-HSPA8 demonstrating preferential recognition of the IDRsTDP-TDP-43 complex by HSPA8. **e,** Schematic illustration of TDP-43 regulation: nuclear autoregulation via exitron splicing generates IDRsTDP, which form complexes with TDP-43 in the cytoplasm, leading to CMA-mediated degradation. **f**-**i,** Analysis of TDP-43 Tg mice injected with AAV-IDRsTDP, including Western blotting (**g**), heatmap and cluster analysis of splicing abnormalities shown in TDP-43 Q331K knock-in mice (**h**) and the effect of IDRsTDP treatment on TDP-43 Tg mice (**i**). Two-way ANOVA (a,c). ANOVA followed by Tukey’s test (**g**). All graphs show mean ± SEM.

Given the presence of a CMA recognition motif within TDP-43^38^, we hypothesized that the TDP-43-IDRsTDP heterocomplex might be targeted for degradation via chaperone-mediated autophagy (CMA). Consistent with this, treatment with the autophagy inhibitor bafilomycin A1 in SH-SY5Y cells led to an increase in endogenous IDRsTDP levels, particularly within the cytoplasm (Fig. 4b, Extended Data Fig. 5b). This finding strongly suggests the involvement of autophagy in regulating the levels of IDRsTDP and the TDP-43-IDRsTDP complex.

To confirm the role of CMA in the degradation of the IDRsTDP-TDP-43 complex, we investigated the effect of HSPA8 depletion, a key CMA component^39^, in HEK293T cells expressing both IDRsTDP and EGFP-tagged TDP-43. HSPA8 knockdown led to an accumulation of exogenously expressed IDRsTDP, supporting its degradation through the CMA pathway (Fig. 4c). Importantly, the effect of HSPA8 depletion on EGFP-TDP-43 levels was dependent on the presence of IDRsTDP. Co-expression with IDRsTDP, but not with IDRsTDP-dN, resulted in increased EGFP-TDP-43 levels upon HSPA8 knockdown, highlighting the essential role of IDRsTDP in mediating HSPA8-dependent regulation of TDP-43 (Fig. 4c). Further supporting a direct interaction between HSPA8 and the IDRsTDP-TDP-43 complex, co-immunoprecipitation experiments revealed that HSPA8 preferentially interacted with IDRsTDP over full-length TDP-43. Moreover, the presence of IDRsTDP enhanced the interaction between HSPA8 and TDP-43 (Fig. 4d). Taken together, these findings strongly suggest that the IDRsTDP-TDP-43 complex is specifically targeted for degradation via a CMA pathway mediated by HSPA8.

Our results unveil a sophisticated, two-pronged mechanism for regulating TDP-43 protein levels (Fig. 4e). Within the nucleus, TDP-43 autoregulates its own expression at the RNA level by promoting exitron splicing, which generates IDRsTDP^7,9^. These IDRsTDP molecules then interact with TDP-43 in the cytoplasm, leading to the formation of hetero-oligomers that are recognized by HSPA8 and targeted for degradation via CMA. This elegant interplay between RNA-level autoregulation and protein-level fine-tuning ensures tight control over TDP-43 levels within the cell.

### IDRsTDP mitigates TDP-43-induced RNA processing abnormalities and neurotoxicity in vivo

To validate the physiological relevance of IDRsTDP-mediated TDP-43 regulation, we employed a transgenic mouse model overexpressing human TDP-43 (C57BL/6-Tg(Prnp-*TARDBP*)3cPtrc/J)^40^. Neonatal mice were intracerebroventricularly injected with AAV-PHP.eB encoding IDRsTDP (Fig. 4f). Analysis of the mouse frontal cortex at 3 months post-injection revealed that IDRsTDP expression effectively reduced the elevated levels of TDP-43 (Fig. 4g), confirming our findings from cellular models.

Given the well-established role of TDP-43 dysregulation in RNA metabolism, we next investigated the impact of IDRsTDP on these processes. Remarkably, IDRsTDP expression ameliorated the transcriptional abnormalities observed in TDP-43 overexpressing mice, shifting their transcriptional profiles closer to those of wild-type mice (Extended Data Fig. 6a, b). Notably, IDRsTDP expression in TDP-43 hemizygous Tg mice significantly rescued splicing defects, specifically those identified in TDP-43 Q331K knock-in mice, a model harboring an ALS-causing mutation (Fig. 4h, i, Extended Data Fig. 6c, d)^21^. Importantly, this beneficial effect on RNA metabolism persisted in a long-term cohort even 7 months post-injection, despite limited detectable IDRsTDP expression at this time point (Extended Data Fig. 6e-h), suggesting a durable impact on RNA processing.

We then examined whether the observed improvements in RNA metabolism translated to functional benefits. TDP-43 homozygous Tg mice exhibit a shortened lifespan^40^. Strikingly, IDRsTDP expression significantly improved survival in female mice (Extended Data Fig. 7a). Furthermore, in a subset of IDRsTDP-treated mice surviving up to approximately 400 days, we observed reduced neuronal loss in the frontal cortex compared to untreated mice (Extended Data Fig. 7b-d), indicating a neuroprotective effect of IDRsTDP.

### Promotion of exitron splicing mitigates TDP-43 pathology caused by proteasome defects

To further investigate the therapeutic potential of enhancing *TARDBP* exitron splicing, we evaluated the effects of promoting this splicing event in vivo. We utilized AS5, an antisense oligonucleotide (ASO) previously validated in human cells, harboring an identical sequence in mice, designed to inhibit HNRNPA1 binding (Extended Data Fig. 8a, b). Intracerebroventricular injection of AS5 into wild-type neonatal mice resulted in robust enhancement of exitron splicing in the cerebrum, brainstem, and spinal cord (Extended Data Fig. 8c-e). Importantly, AS5 exhibited high specificity, promoting only exitron splicing without affecting overall *TARDBP* expression levels or intron 7 splicing (Extended Data Fig. 8f, g). Notably, this effect was long-lasting, persisting in the spinal cord for at least 12 weeks, accompanied by a modest decrease in TDP-43 protein levels (Extended Data Fig. 8i, j). Intrathecal administration in adult mice also confirmed this splicing-promoting effect, with no adverse effects on body weight, grip strength, or inflammation markers (Extended Data Fig. 8h, k-o).

To assess the therapeutic potential of exitron splicing modulation in a TDP-43 proteinopathy model independent of TDP-43 overexpression, we employed *Rpt3*^flox/flox^;*VAChT-Cre*^+/-^ mice. These mice exhibit motor neuron-specific proteasome deficiency, leading to TDP-43 accumulation and recapitulating key aspects of TDP-43 proteinopathies^41^. By eight weeks of age, these mice display characteristic pathological features, including increased cytoplasmic TDP-43, reduced nuclear TDP-43, and the presence of TDP-43 aggregates in a subset of spinal motor neurons (Extended Data Fig. 9a, b). Additionally, we observed progressive motor neuron death, evidenced by ubiquitin-positive fragmented neurons surrounded by activated microglia (Extended Data Fig. 9c-e).

Remarkably, AS5 administration to neonatal *Rpt3*^flox/flox^;*VAChT-Cre*^+/-^ mice resulted in a significant increase in surviving motor neurons and a reduction in TDP-43 aggregation at eight weeks of age, correlating with the efficiency of exitron splicing (Fig. 5a-c). Moreover, long-term AS5 treatment (intracerebroventricularly at birth, followed by intrathecal administration at 16 and 30 weeks) significantly extended survival compared to untreated mice (Fig. 5d).

**Fig.5.**
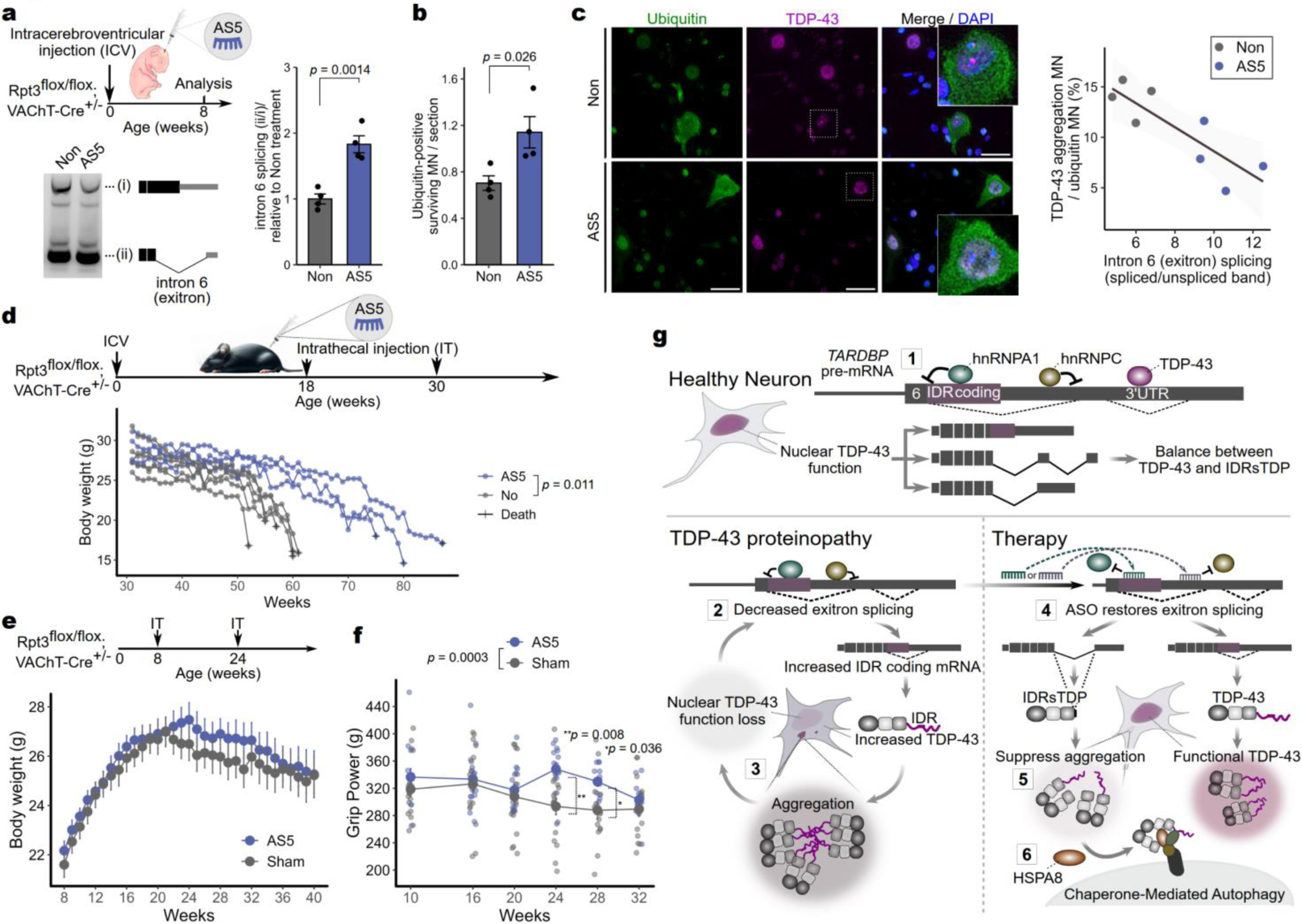
Splicing-promoting effect of *TARDBP* exitron in motoneuron-specific *Rpt3* knockout mice. **a-c,** Effect of AS5 treatment on exitron splicing (**a),** motoneuron viability (**b**), and TDP-43 aggregation (**c**) in Rpt3^flox/flox^;VAChT-Cre^+/-^ mice. Scale bar: 30 μm. **d,** Survival analysis of Rpt3^flox/flox^;VAChT-Cre^+/-^ mice treated with AS5. **e,f,** Body weight (**h**) and grip strength measurements (**f**) of AS5-treated and sham surgery groups. **g,** Graphical abstract on molecular mechanisms of therapeutic strategies targeting *TARDBP* exitron splicing. Data are presented as mean ± SEM. Statistical tests: Student’s t-test (**a,b**), log-rank test (**d**), repeated measures ANOVA with post-hoc tests (**f**).

Importantly, even when AS5 treatment was initiated after the onset of TDP-43 accumulation (intrathecal administration at 8 and 24 weeks), it effectively attenuated the decline in grip strength, independent of body weight loss (Fig. 5e, f; Extended Data Fig. 9f, g), although it did not significantly alter the final survival duration (Extended Data Fig. 9h).

These results demonstrate the efficacy of promoting *TARDBP* exitron splicing as a therapeutic strategy for TDP-43 proteinopathies caused by proteasomal abnormalities, highlighting its potential to mitigate disease progression.

## Discussion

This study unveils a novel therapeutic strategy for TDP-43 proteinopathies by targeting the intrinsic autoregulatory mechanism governing TDP-43 homeostasis. Central to this mechanism is the splicing of an exitron within *TARDBP* pre-mRNA, which encodes the aggregation-prone IDRs of TDP-43 (Fig. 5g-1).

In TDP-43 proteinopathies, we observe a significant reduction in exitron splicing due to impaired TDP-43 nuclear function (Fig. 5g-2), exacerbating a pathological cascade characterized by increased TDP-43 aggregation and diminished nuclear TDP-43 function (Fig. 5g-3). Our work identifies HNRNPA1 and HNRNPC as key repressors of this exitron splicing event. We demonstrate that ASOs targeting their binding sites effectively restore exitron splicing (Fig. 5g-4), thereby interrupting the pathological cycle. This intervention simultaneously suppresses the production of aggregation-prone IDR-containing TDP-43 isoforms and increases the levels of the protective IDR-spliced-out TDP-43 isoforms (IDRsTDP).

Our findings suggest that IDRsTDP plays a multifaceted role in TDP-43 regulation. While excessive IDRsTDP levels can lead to cellular toxicity^10^, likely through excessive sequestration of endogenous TDP-43 via the NTD, our data support a model where tightly regulated IDRsTDP levels are beneficial. By interacting with TDP-43, IDRsTDP appears to modulate liquid-liquid phase separation, a process crucial for TDP-43 function^22^, and targets the resulting complex for degradation via CMA. Importantly, although AAV-mediated IDRsTDP overexpression in wild-type mice did not induce overt toxicity, therapeutic strategies directly targeting IDRsTDP expression warrant careful consideration to avoid perturbing normal TDP-43 function in healthy cells.

Physiologically, IDRsTDP levels are finely tuned through a combination of TDP-43-mediated autoregulatory exitron splicing and nonsense-mediated decay^7,9^, allowing a controlled interaction with excess TDP-43 to mitigate aggregation. This protein-level regulation provides a rapid response mechanism that complements the slower RNA-level control exerted through exitron splicing. In TDP-43 proteinopathies, however, the dysregulation of nuclear TDP-43 disrupts this balance, leading to increased expression of IDR-containing TDP-43 isoforms, reduced IDRsTDP production, and ultimately impaired aggregation suppression and CMA-mediated degradation (Fig. 5g-5, 6).

While our study highlights the promise of modulating exitron splicing as a therapeutic strategy, further validation is necessary to realize its full translational potential. One key consideration is the delivery of ASOs to the CNS in humans. Intrathecal administration, which has shown promise for other neurological disorders, or the use of modified ASO chemistries with enhanced CNS penetration could be explored^42,43^. Additionally, while our ASOs were highly specific in our models, it will be important to evaluate potential off-target effects on global splicing patterns in future studies. To enhance efficacy and safety, further optimization of ASO design through a deeper understanding of HNRNPA1 and HNRNPC’s regulatory roles will be essential. Moreover, it will be crucial to determine whether enhancing exitron splicing is effective in models bearing ALS/FTLD-causing mutations, which are known to impair TDP-43 autoregulation^20,21^, and in other TDP-43 proteinopathies beyond those investigated here. Furthermore, exploring the role of age-related factors, such as DNA demethylation, in exitron splicing dysregulation^44^, is warranted. Addressing these questions will pave the way for translating our findings into much-needed therapies for these devastating diseases.

Our approach represents a paradigm shift from conventional strategies that often target downstream consequences or aim to globally suppress protein expression. By leveraging the intrinsic autoregulatory nature of *TARDBP* exitron splicing, which serves as a sensitive barometer of TDP-43 function, we directly address the core pathogenic mechanisms underlying TDP-43 proteinopathies. As we advance this strategy towards clinical application, we envision a future where we can offer new hope and tangible therapeutic options to patients suffering from ALS, FTLD, and other TDP-43-related disorders.

## Acknowledgements

This work was supported by JSPS KAKENHI Grant Numbers JP20K07880 and JP17K09751, AMED under Grant Number JP24ek0109603h0003 and JP21m0203001j0005, Yujin Memorial Grant (Niigata University School of Medicine), Tsubaki Memorial Foundation, Kyowa-kai medical research Fellowship (Niigata University School of Medicine), and SERIKA FUND. We are grateful to Yoshitaka Tashiro (National Center for Geriatrics and Gerontology) and Ryosuke Takahashi (Kyoto University) for providing the *Rpt3*tm1 mice through the Center for Animal Resources and Development, Kumamoto University, and for their valuable advice. Both were affiliated with Kyoto University when they originally developed and published these mice^41^. We thank Masato Yano (Graduate School of Medical and Dental Sciences, Niigata University) for critically reading the manuscript and providing valuable comments.

## Authors and Affiliations

Department of Neurology, Clinical Neuroscience Branch, Brain Research Institute, Niigata University, Niigata, Japan: Takuma Yamagishi, Shingo Koide, Genri Toyama, Aya Washida, Yumi Yamada, Ryutaro Hanyu, Ekaterina Nadbitova, Takuya Konno, Tomohiko Ishihara, Osamu Onodera, Akihiro Sugai Department of Molecular Neuroscience, Resource Branch for Brain Disease Research, Brain Research Institute, Niigata University, Niigata, Japan: Yuka Mitsuhashi Koike, Taisuke Kato, Osamu Onodera.

Contributions: A.S. designed the research concept. A.S. and T.Y. designed experiments for IDRsTDP. A.S., A.W., T.KN., and T.KT. performed ASO experiments. T.Y., Y.Y., R.H., A.W., G.T., and A.S. conducted in vitro IDRsTDP experiments. T.Y., A.W., S.K., and E.N. carried out experiments on TDP-43 Tg mice. A.S. analyzed and interpreted RNA-seq data. T.Y., A.S., Y.M.K., T.I., and O.O. interpreted data. A.S. and T.Y. wrote the manuscript with significant contributions from O.O. and input from all other authors.

Corresponding author: Akihiro Sugai.

## Methods

### Analysis of RNA-seq data from the NYGC ALS Consortium

We analyzed RNA-seq data from the NYGC ALS dataset (GSE153960), including frontal cortex (40 healthy, 122 ALS-TDP, 24 FTLD-TDP, 23 ALS-FTLD) and cerebellum (27 healthy, 135 ALS-TDP, 39 FTLD-TDP, 12 ALS-FTLD) samples. Cases were excluded based on NYGC ALS Consortium metadata: RNA integrity number (RIN) ≤ 5, SOD1 mutations, FTD with FUS or Tau aggregates, and ALS with suspected Alzheimer’s disease (Fig. 1b). Additionally, cases with unknown or imprecise age at death were excluded from the analysis.

Raw fastq files were quality-trimmed using fastp v0.20.0^45^ and aligned to GENCODE Release 39 (GRCh38.p13) using STAR v2.7.3a^46^ with default parameters. Multiple BAM files from the same subject and tissue were merged using samtools v1.7^47^. Splicing site usage was analyzed using McSplicer v2.1.0^48^ with GENCODE Release 39 as the reference annotation. Acceptor site usage rates for *TARDBP* intron 6 (exitron) and intron 7 were calculated using 100 bootstrap iterations, with the median value used for analysis. Total splicing rates were computed as: 3’-side acceptor utilization + 5’ side acceptor utilization × (1 - 3’-side acceptor utilization).

*STMN2* expression data were obtained from NYGC ALS Consortium metadata. Multiple regression analysis was performed on log-transformed *STMN2* TPM values using R v4.4.0. Variables included sequencing platform, RIN, age at death, sex, brain region, C9orf72 repeat expansion status, TDP-43 proteinopathy status, and log-transformed *TARDBP* intron 6 splicing rates. All variables were standardized before analysis. The full model was constructed using the lm() function, and model selection was performed using the dredge() function (MuMIn package) with AIC and BIC criteria. The best AIC model was selected for final interpretation. Standardized partial regression coefficients and 95% confidence intervals were visualized using ggplot2 v3.5.1. Model fit was assessed using the adjusted R-squared value and overall model significance.

### Analysis of ENCORE project shRNA RNA-seq data

RNA-seq data from the ENCODE project’s shRNA experiments in HepG2 cells were analyzed using BAM files aligned to the GRCh38 genome assembly with annotation version 29. McSplicer v2.1.0^48^ was employed to quantify *TARDBP* exitron acceptor site usage, following the same methodology described for the NYGC ALS Consortium data analysis. Results were compared to corresponding control experiments. ENCODE eCLIP data for HNRNPA1 (ENCFF706CRY) and HNRNPC (ENCFF774JCA) binding profiles are presented.

### Evaluating the effects of ASOs on exitron splicing enhancement

HEK293T cells were cultured in DMEM with 10% FBS. 10 μM Morpholino ASOs (GeneTools) were delivered using 6 μM Endo-Porter PEG (GeneTools). ASOs targeting HNRNPA1 (AS2-AS6.2) and HNRNPC (CS1-CS4) binding regions of the *TARDBP* exitron were designed. Standard Control Morpholino Oligo (Ctrl) served as a negative control. RNA was extracted 30 hours post-treatment. For sensitive splicing detection, 100 μg/ml cycloheximide (Wako, 037-20991) was added 6 hours before harvest. iPSC-derived neurons (ReproNeuro, RCDN001N) were treated with ASOs 14 days post-seeding. For TDP-43 mislocalization studies, HEK293T cells were transfected with CSE1L siRNA (L-004413-00-0005, 20 nM) using Lipofectamine RNAiMAX (Thermo Fisher) before ASO treatment. RNA was isolated using NucleoSpin RNA (Takara, 740955) and reverse-transcribed with PrimeScript RT Master Mix (Takara, RR036A). The exitron region was amplified using LA Taq (Takara, RR002A). PCR products were analyzed on 2% agarose gels and imaged using an Amersham Imager 680 (GE Healthcare).

### Prediction of RNA binding protein binding to exitron using RBPNet

RBPNet v0.0.2^27^ was used to predict HNRNPA1 and HNRNPC binding to the *TARDBP* exitron. Control and modified sequences (with 25 or 30 bases replaced by “N”) were analyzed. To quantify differences in binding profiles, Kullback-Leibler divergence was calculated using R.

### Generation of *TARDBP* minigene and splicing analysis

A *TARDBP* minigene (exon 5 to 6) was cloned into pCDNA3.1/myc-His(-) A. HEK293T cells were transfected with HNRNPA1 (Ref#106750884) or HNRNPC (Ref#106585366) DsiRNAs. Knockdown was assessed by Western blotting using anti-hnRNPA1 (Cell Signaling, 4296) and anti-hnRNP C1/C2 (SCB, sc-32308) antibodies. Exitron splicing was analyzed by RT-PCR.

### *STMN2* expression and exitron splicing in SH-SY5Y cells

SH-SY5Y cells were cultured in DMEM/F12 GlutaMax medium (Thermo Fisher, 10565018) with 10% FBS and were transfected with CSE1L siRNA (L-004413-00-0005, 20 nM) using Lipofectamine RNAiMAX followed by ASO delivery 24 hours later. RNA was extracted 48 hours post-ASO treatment. Quantitative real-time PCR was performed using SYBR Green Premix ExTaq II on a Thermal Cycler Dice Real Time System III (Takara Bio).

### TDP-43 nuclear-to-cytoplasmic ratio analysis

SH-SY5Y cells were seeded on μ-Slide 8 well ibiTreat slides (ibidi, ib80826) coated with Matrigel (Corning, 356231) diluted 6-fold in Opti-MEM™ I (Thermo Fisher, 31985062). Cells were cultured in DMEM/F12 GlutaMax with 10% FBS. The day after seeding, CSE1L knockdown was performed. Twenty-four hours post-knockdown, the medium was replaced with DMEM/F12 GlutaMax containing 1% FBS, 10 μM all-trans-Retinoic Acid (FUJIFILM Wako, 186-01114), and 50 ng/ml BDNF (FUJIFILM Wako, 028-16451), and ASOs were simultaneously delivered.

IPS cell-derived neurons (ReproNeuro, ReproCell, RCDN001N) were seeded on μ-Slide 8 well ibiTreat slides coated according to the manufacturer’s protocol and cultured in 200 μl of the provided medium. On day 13 post-seeding, CSE1L knockdown was performed using a recombinant lentivirus (LVS(VB230809-1203tuz)-K1) containing shRNA targeting hCSE1L (pLV[shRNA]-Puro-U6>hCSE1L[shRNA#1]) at an MOI of 5 in 150 μl of medium. A control lentivirus (LVS(VB010000-9460fht)-K1) with scrambled shRNA (pLV[shRNA]-Puro-U6>Scramble[shRNA#1]) was used at the same MOI. ASOs were delivered on day 14.

For immunocytochemistry, SH-SY5Y cells were fixed 72 hours post-ASO delivery, and iPS cell-derived neurons on day 20 (6 days post-ASO delivery). Cells were fixed with 4% PFA, permeabilized with 0.1% Triton X-100, and blocked with 5% BSA. Primary antibodies (rabbit anti-TDP-43, 1:200, Proteintech 12892-1-AP; mouse anti-TUJ1, 1:400, Abcam, ab7751) were applied overnight at 4°C, followed by secondary antibodies (Alexa Fluor 568 goat anti-rabbit IgG and Alexa Fluor 488 goat anti-mouse IgG, both 1:400) for 1 hour. Cells were mounted with DAPI-containing medium (ibidi, ib50011). Images were acquired using a BIOREVO BZ-9000 microscope (Keyence). Nuclear and cytoplasmic ROIs were defined using ImageJ based on DAPI and TUJ1 signals. TDP-43 fluorescence intensity was measured in >200 cells per condition for SH-SY5Y cells and >400 neurons per condition for iPS cell-derived neurons. For statistical analysis, 200 SH-SY5Y cells or 400 iPS cell-derived neurons were randomly selected for each condition using R.

### Insoluble TDP-43 analysis after CSE1L knockdown and ASO transduction

SH-SY5Y cells were seeded in 6-well plates coated with Matrigel. After CSE1L knockdown and ASO delivery as described above, cells were harvested at 52 hours. Cells were lysed with RIPA buffer (NACALI, 16488-34) containing protease inhibitor cocktail (Sigma, P8340). Lysates were sonicated and centrifuged at 20,000 × g for 30 min at 4°C. The supernatant (RIPA-soluble fraction) was collected. The pellet was treated with urea buffer (7 M urea, 2 M thiourea, 4% CHAPS, 30 mM Tris pH 8.5, protease inhibitors), sonicated, and centrifuged at 20,000 × g for 30 min at 22°C. This supernatant formed the urea-soluble fraction. Protein concentrations were determined using Pierce BCA Protein Assay Kit (Thermo Scientific, 23225). Samples were subjected to SDS-PAGE and Western blotting using anti-TDP-43 antibody (Proteintech, 12892-1-AP) and HRP-conjugated secondary antibody (Dako, P0448). Total protein was visualized using No-Stain Protein Labeling (Thermo Fisher, A44449).

### Nuclear and cytoplasmic TDP-43 levels upon CSE1L knockdown

SH-SY5Y cells were subjected to CSE1L knockdown for 72 hours. Nuclear and cytoplasmic proteins were extracted using the NE-PER Nuclear and Cytoplasmic Extraction Reagents (Thermo Scientific, 78833). For Western blotting, primary antibodies against TDP-43 (Proteintech, 12892-1-AP), CSE1L (Abcam, ab151546), GAPDH (MBL, M171-3), and Lamin A/C (Cell Signaling, 4777), followed by appropriate HRP-conjugated secondary antibodies (Goat anti-Rabbit IgG or Goat anti-Mouse IgG) were used.

### Plasmid construction and TDP-43 protein purification

A plasmid encoding human TDP-43 with a C-terminal His×6-MBP tag was purchased from Addgene (pJ4MTDP-43, Addgene #104480). Two additional plasmids encoding IDRsTDP-2(Extended Data Fig. 4a) (VB210204-1240sfb) and IDRsTDP-dN (VB210422-1361jgv) were constructed by VectorBuilder. A TDP-ex5 encoding plasmid was generated by introducing a deletion in pJ4MTDP-43 using the KOD-Plus-Mutagenesis Kit (TOYOBO, SMK-101).

Protein expression and purification were performed as previously described^49,50^, with minor modifications. Briefly, Rosetta2(DE3)pLysS cells were used for protein expression. After IPTG induction and overnight incubation, cells were lysed, and proteins were purified using a HisTrap HP column followed by an Amylose Resin column. Purified proteins were snap-frozen and stored at −80°C.

### Observation and quantification of liquid-liquid phase separation

After buffer exchange (20 mM HEPES-NaOH (pH 7.4), 150 mM NaCl, and 1 mM DTT), full-length TDP-43 and IDRsTDP or IDRsTDP-dN proteins were mixed and diluted to a final concentration of 5 μM each. TEV protease was added and the mixture was incubated for 30 minutes. Following incubation, a portion of the sample was placed on a glass slide, covered with a coverslip, and the droplets formed by LLPS were observed and imaged using differential interference contrast (DIC) microscopy (IX83, Olympus). The number and area of droplets in the images were measured using the Trainable Weka Segmentation plugin in Fiji.

### Aggregation studies

Purified recombinant proteins (TDP-43, IDRsTDP, IDRsTDP-dN, and TDP-ex5) were thawed on ice, concentrated using Amicon Ultra Centrifugal Filters with a 10 kDa MWCO (Millipore), and buffer-exchanged into 20 mM HEPES-NaOH (pH 7.4), 150 mM NaCl, and 1 mM DTT using Zeba Spin Desalting Columns with a 7K MWCO (Thermo Scientific). Protein concentrations were determined using the Pierce BCA Protein Assay Kit-Reducing Agent Compatible (Thermo Scientific). Proteins were diluted in buffer (20 mM HEPES (pH 7.0), 150 mM NaCl, and 1 mM DTT) to a final concentration of 3 μM for each protein. Aggregation was monitored by measuring turbidity at 405 nm using a FilterMax F5 microplate reader (Molecular Devices). The assay was initiated by adding TEV protease (New England Biolabs) to cleave the MBP tag from the TDP-43-MBP constructs. Turbidity was measured every 5 minutes while shaking at 30°C. Values were expressed as the change in turbidity from the starting point and normalized to 100% based on the average change in TDP-43 alone or TDP-43+IDRsTDP-dN at the endpoint.

### Observation and quantification of aggregates

Following the aggregation assay, the aggregated samples were dispersed on carbon film supported copper grids and negatively stained with 2% (w/v) phosphotungstic acid solution (pH 7.0) for 15 seconds. Samples were imaged using a transmission electron microscope (JEM-1400Plus; JEOL Ltd., Tokyo, Japan). Sample staining and imaging were performed by Tokai Electron Microscopy, Inc. The area of the aggregates in the images was measured using the Trainable Weka Segmentation plugin in Fiji.

### Split-GFP complementation assay

HEK293T cells were transfected with plasmids encoding the Split-GFP system components using X-tremeGENE HP DNA Transfection Reagent (Roche, 06366236001). Two plasmids were compared in this experiment: TriGFP + IDRsTDP-myc (VB220113-1341hvf) and TriGFP + IDRsTDP-dN-myc (VB220113-1318frz). Two days post-transfection, cells were subjected to osmotic stress by adding sorbitol to the medium (final concentration 0.4 M) for 1 hour. For immunocytochemistry, GFP-Booster (1:500, ChromoTek, GB2AF48850) and Anti-Myc (1:200, Abcam, ab32) were used and Alexa Fluor 568 goat anti-mouse IgG (1:500) secondary antibody was applied. Immunoprecipitation was performed using the GFP-Trap Magnetic Agarose kit (Proteintech, GTMAK-20) following the manufacturer’s instructions using RIPA buffer with protease inhibitor cocktail. Input and eluted fractions were analyzed by SDS-PAGE and Western blot using a mouse monoclonal antibody against TDP-43 (2E2-D3, Abcam, ab57105) and a goat anti-mouse IgG secondary antibody.

### TDP-43 stability assay

HEK293T cells were transfected with either IDRsTDP-myc (VB210901-1048nsv) or IDRsTDP-dN-myc (VB220106-1552mwb) plasmids. At 48 hours post-transfection, cells were treated with 25 μg/ml cycloheximide and harvested at 0, 6, and 12 hours post-treatment. Proteins were extracted using RIPA buffer and analyzed by Western blotting. For detection, primary antibodies against TDP-43 (Proteintech 10782-2-AP, Fig.4a; Cosmo Bio TIP-TD-P09, Extended Fig.5a) and HRP-conjugated polyclonal goat anti-rabbit antibody were used. Endogenous TDP-43 stability was evaluated by comparing TDP-43 protein levels relative to total protein at each time point after CHX treatment in cells expressing either IDRsTDP-dN or IDRsTDP. The interaction between transfection condition and CHX treatment duration on TDP-43 degradation was analyzed using two-way ANOVA.

### Bafilomycin A1 treatment in SH-SY5Y cells

SH-SY5Y cells were cultured in DMEM/F12 GlutaMax with 10% FBS. Cells were treated with either DMSO (Sigma-Aldrich, D2650; final concentration 0.01%) or bafilomycin A1 (Sigma-Aldrich, 19-148; 10 nM, dissolved in DMSO) for 48 hours. Proteins were extracted using RIPA buffer or NE-PER Nuclear and Cytoplasmic Extraction Reagents containing protease inhibitors. For Western blotting, primary antibodies against TDP-43 (Proteintech 10782-2-AP, Cosmo Bio TIP-TD-P09) were used, followed by HRP-conjugated polyclonal goat anti-rabbit antibody.

### HSPA8 knockdown assay

HEK293T cells were transfected with 20 nM siRNA (siHSPA8 or negative control, riFECTa Kit DsiRNA Duplex, Integrated DNA Technologies) using Lipofectamine RNAiMAX. The following day, the medium was replaced, and cells were transfected with plasmids expressing IDRsTDP-myc (VB210901-1048nsv) or IDRsTDP-dN-myc (VB220106-1552mwb), and EGFP-TDP-43 (VB240423-1225beg) and EGFP (transfection control) using Lipofectamine 3000. At 72 hours post-transfection, cells were lysed with RIPA buffer containing protease inhibitors. For Western blotting, primary antibodies against TDP-43 (Proteintech, 10782-2-AP), GFP (Proteintech, 50430-2-AP), and HSPA8 (Santa Cruz, sc-7298) were used, followed by HRP-conjugated secondary antibodies (Dako, P0448 and P0447).

### EGFP-HSPA8 co-immunoprecipitation assay

HEK293T cells were transfected with combinations of EGFP-HSPA8(VB240309-1040zsb), TDP-43, and IDRsTDP-myc (VB210901-1048nsv) or IDRsTDP-dN-myc (VB220106-1552mwb) plasmids using X-tremeGENE™ HP DNA Transfection Reagent. The TDP-43 expression plasmid was generated by inserting human TDP-43 cDNA into pcDNA DEST40 vector (Invitrogen). Thirty-six hours post-transfection, cells were subjected to co-immunoprecipitation using the GFP-Trap Magnetic Agarose Kit following the manufacturer’s instructions. Input and co-immunoprecipitated samples were analyzed by Western blotting using antibodies against TDP-43 (Proteintech, 10782-2-AP) and HSPA8 (Santa Cruz, sc-7298), followed by HRP-conjugated secondary antibodies.

### TDP-43 transgenic mice

All procedures were approved by the Institutional Animal Care and Use Committee of Niigata University. C57BL/6-Tg(Prnp-TARDBP)3cPtrc/J (JAX stock #016608) mice were used. TDP-43Tg/+ mice were maintained by crossing with wild-type mice. Standard housing conditions were provided. Humane endpoints were applied as approved in the animal experiment protocol, euthanizing mice when they became unable to move, showed a rapid decrease in body weight, or a reduction of more than 25% from their maximum weight.

Offspring from hemizygous crosses were used for the study. AAV administration was performed on postnatal day 1. Treatment was randomly assigned, with PBS administered to the control group. Blinding was ensured through PCR detection of vector sequences at autopsy or death. Mice were weaned at 3 weeks of age. Homozygous mice were housed separately and provided with Diet Gel 76A (ClearH2O) on the cage floor. Forelimb grip strength was measured using a grip strength meter (GPM-100V, Melquest). Each measurement was performed thrice, and the maximum value was used for analysis. Mice were dissected at predetermined timepoints or study endpoints.

Genotyping was performed using ddPCR for human TARDBP quantification in offspring from hemizygous pairs, and PCR for human TARDBP exons 5-6 in offspring from hemizygous-wildtype crosses. Treatment groups were identified by PCR detection of vector genes in brain tissue.

### Administration of AAV-IDRsTDP to mice

AAVs expressing IDRsTDP-2 (Extended Data Fig. 4a), ssAAV-PHP.eB-hSYN-IDRsTDP (6.53×10^13 GC/ml in PBS) and ssAAV-PHP.eB-CMV-IDRsTDP-myc (3.97×10^13 GC/ml in PBS), were designed and packaged using VectorBuilder’s service. Intracerebroventricular injection of AAVs was performed following a previously described protocol^51^. P1 neonatal mice were cryo-anesthetized on ice and a glass capillary needle was prepared, filled with 3 μl of either: ssAAV-PHP.eB-SYN-IDRsTDP diluted 1.5-fold in PBS (total 1.31×10^11 GC/body), undiluted ssAAV-PHP.eB-CMV-IDRsTDP-myc (total 1.19×10^11 GC/body), or PBS (control). The needle was attached to a polyethylene tube connected to a 25 μl Hamilton microsyringe. The needle was inserted into the brain of the immobilized mouse at a point 2 mm to the right and 3 mm rostral from the bregma. The AAV solution was administered using a syringe pump at a rate of 1.1 μl/sec. After confirming complete administration without resistance, the needle was withdrawn. The absence of prolonged bleeding was verified, the mice were warmed, and returned to their cages.

### Western blotting of TDP-43-transgenic mouse brain tissue

Mouse frontal cortex tissues were homogenized in RIPA buffer with protease inhibitors, sonicated, and ultracentrifuged at 100,000 × g for 30 minutes at 4°C. Protein samples were analyzed by Western blotting using anti-TDP-43 (Proteintech 10782-2-AP) and HRP-conjugated goat anti-rabbit IgG antibodies.

### RNA analysis of TDP-43-transgenic and Q331K knock-in mice

RNA from frontal cortex was assessed for concentration and integrity (RIN) using RNA Screen Tape (Agilent, 5067-5576) on an Agilent 2200 TapeStation. cDNA was synthesized using PrimeScript™ RT Master Mix. RNA sequencing was performed by Genome-Lead Co., Ltd. (Kagawa, Japan) using KAPA mRNA Capture Kit (KAPA, KK8440) and MGIEasy RNA Directional Library Prep Set (MGI, 1000006385) on a DNBSEQ-T7RS platform (MGI).

Sequencing reads were processed using fastp v0.20.0^45^, aligned to the mouse reference genome (GRCm39/mm39) using STAR v2.7.3a^46^, and quantified with TPMCalculator v0.0.3^52^. Differential transcript analysis was performed using DESeq2 v1.40.2^53^ in R v4.4.0, comparing TDP-43 transgenic mice to wild-type controls. Heat maps and principal component analysis were performed using pheatmap and FactoMineR package v2.8^54^, respectively.

Splicing analysis utilized McSplicer v2.1.0.^48^ For TDP-43 Q331K knock-in mice (RJNA388188), the reference GTF file (gencode.vM28.primary_assembly.annotation.gtf) was modified to exclude seven genes (Ccl27a, Zfp637, Arrb1, Med16, Rpl38, Mirg, Wdr45) that caused McSplicer errors. Differential splicing was assessed using the false discovery rate (FDR) method. For TDP-43-transgenic mice, analysis focused on genes showing significant splicing changes (FDR < 0.05, excluding Tardbp) in the Q331K knock-in analysis.

### Immunostaining of TDP-43-transgenic mouse brain tissue

Cortical tissue from TDP-43 transgenic mice was fixed, paraffin-embedded, and sectioned at 4 μm thickness using a microtome (REM-710, YAMATO). Sections were mounted on MAS-coated glass slides (MATSUNAMI) and subjected to deparaffinization and antigen retrieval in Citrate Buffer (pH 6.0; Abcam, ab64236). Following blocking with 5% bovine serum albumin and permeabilization with 0.1% Triton X-100 in phosphate-buffered saline (PBS), sections were incubated with anti-NeuN primary antibody (Merck, MAB377) and Alexa Fluor 488 goat anti-mouse IgG. Autofluorescence was minimized by treatment with TrueBlack(R) Lipofuscin Autofluorescence Quencher (Biotium, 23007). Sections were mounted using VECTASHIELD PLUS Antifade Mounting Medium with DAPI (Vector Laboratories, H-2000-10).

Images were acquired using a BIOREVO BZ-9000 microscope. Neuronal density in the left cortex of 19 homozygous mice (10 AAV-injected, 9 PBS-injected) was analyzed using 10 brain sections per animal. Poorly stained sections were excluded from analysis. Images of the cortical cross-sections were captured, maintaining consistent width. Cells were segmented and counted utilizing Cellpose 2.0 software^55^. Cell density was calculated by dividing the total cell count by the cortical area, measured using Fiji.

### Intracerebroventricular administration of ASO to neonatal mice

Intracerebroventricular (ICV) injection of ASOs was performed on P1 mice as described above for AAV administration, with minor modifications. The glass capillary needle was filled with 2 μl (for survival analysis) or 3 μl (for pathology analysis) of 4 mM morpholino ASO or 1×PBS (control). ASO administration and post-injection care were carried out as described for AAV injection.

### Intrathecal administration of ASO to mice

Intrathecal injection of ASO was performed as previously described^11^. Adult mice were anesthetized and a 32-gauge needle was inserted into the cerebrospinal cavity between the fifth and sixth lumbar vertebrae. A solution of 4 mM morpholino ASO in PBS (15 μl) was injected at a rate of 1 μl/min. The needle was left in place for 15 minutes post-injection before withdrawal. Mice were maintained at an appropriate body temperature throughout the procedure and recovery period.

### Generation and Experimental use of *Rpt3*^flox/flox^;*VAChT-Cre*^+/-^ mice

All experimental procedures were conducted in accordance with protocols approved by the Institutional Animal Care and Use Committee of Niigata University. *Rpt3*tm1 mice (floxed *Rpt3*; CARD ID 1851) were obtained from the Center for Animal Resources and Development (CARD), Kumamoto University, and maintained as homozygotes. These were crossed with C57BL/6-Tg(SLC18A3-cre)KMisa mice (*VAChT-Cre*.Early; RBRC01515) from RIKEN BRC to generate *Rpt3*^flox/flox^;*VAChT-Cre*^+/-^ mice. Experimental animals were produced by mating male *Rpt3*^flox/flox^;*VAChT-Cre*^-/-^ with female *Rpt3*^flox/flox^;*VAChT-Cre*^+/-^ mice.

To assess the effects of AS5 treatment, neonatal *Rpt3*^flox/flox^;*VAChT-Cre*^+/-^ mice received ICV injections of AS5. At 8 weeks of age, spinal cords were analyzed for motoneuron viability, TDP-43 aggregation, and exitron splicing (n=4 per group, treated and untreated). For survival analysis, mice were divided into untreated (n=6) and AS5-treated (n=3) groups. The AS5-treated group received ICV injections neonatally and intrathecal (IT) injections at 18 and 30 weeks. Survival was compared using the log-rank test. For phenotype experiments, treatment was randomly assigned to half of the littermates. Mice with no evidence of cerebrospinal fluid reflux during ASO administration were excluded from the analysis. Mice received IT AS5 treatment (n=12) or sham surgery (n=16) at 8 and 24 weeks of age. Body weights were recorded up to 40 weeks, and grip strength was measured at specified time points. Male mice were used for motor function and survival analyses, while both sexes were included in pathological studies. Housing conditions were consistent across all experiments. Mice were euthanized when their body weight decreased by 25% relative to their 4-week weight, as the humane endpoint.

### Immunostaining of spinal cord tissue from *Rpt3*^flox/flox^;*VAChT-Cre*^+/-^ mice

Cervical and thoracic regions of the spinal cord were processed for immunohistochemistry as described for TDP-43 transgenic mouse brain tissue, with some modifications. Serial 4 μm sections were cut and mounted on MAS-coated glass slides, with eight sections per slide at approximately 100 μm intervals. Primary antibodies against TDP-43 (Proteintech, 12892-1-AP), TUJ1 (Abcam, ab7751), ubiquitin (Santa Cruz Biotechnology, SC8017), Rpt3 (Abcam, ab196589), and Iba1 (Wako, 019-19741) were applied overnight at 4°C. Alexa Fluor 488 goat anti-mouse IgG and Alexa Fluor 568 goat anti-rabbit IgG were used as secondary antibodies. Images were captured and processed as described earlier. For quantification, 48 spinal cord sections per mouse were analyzed for surviving ubiquitin-positive motor neurons, and 64 sections for TDP-43 aggregation. Results were expressed as the average number of ubiquitin-positive surviving motor neurons per section and the percentage of TDP-43 aggregation-positive neurons among ubiquitin-positive motor neurons for each mouse.

### Statistics

Statistical analyses were performed based on the nature of the data and experimental design. Normality and homogeneity of variances were assessed using appropriate tests. For two-group comparisons, Student’s t-test or Welch’s t-test was used. Multiple group comparisons employed one-way ANOVA with Tukey’s HSD post-hoc test, Kruskal-Wallis test with Dunn’s post-hoc test, or Welch’s ANOVA with Games-Howell post-hoc test, as appropriate. Dunnett’s test was used for comparing multiple treatment groups against a control. Survival analysis used the Kaplan-Meier method with log-rank test. Longitudinal grip strength data were analyzed using repeated measures ANOVA and ANCOVA. Neuron density in TDP-43-Tg-homo mice was analyzed using one-way ANOVA and a mixed-effects model. Data are presented as mean ± SEM or mean ± SD. All tests were two-sided, with p < 0.05 considered statistically significant.

