## Extended Data Figure for "ASO-enhancement of *TARDBP* exitron splicing mitigates TDP-43 proteinopathies"

#### Extended Data Fig.1

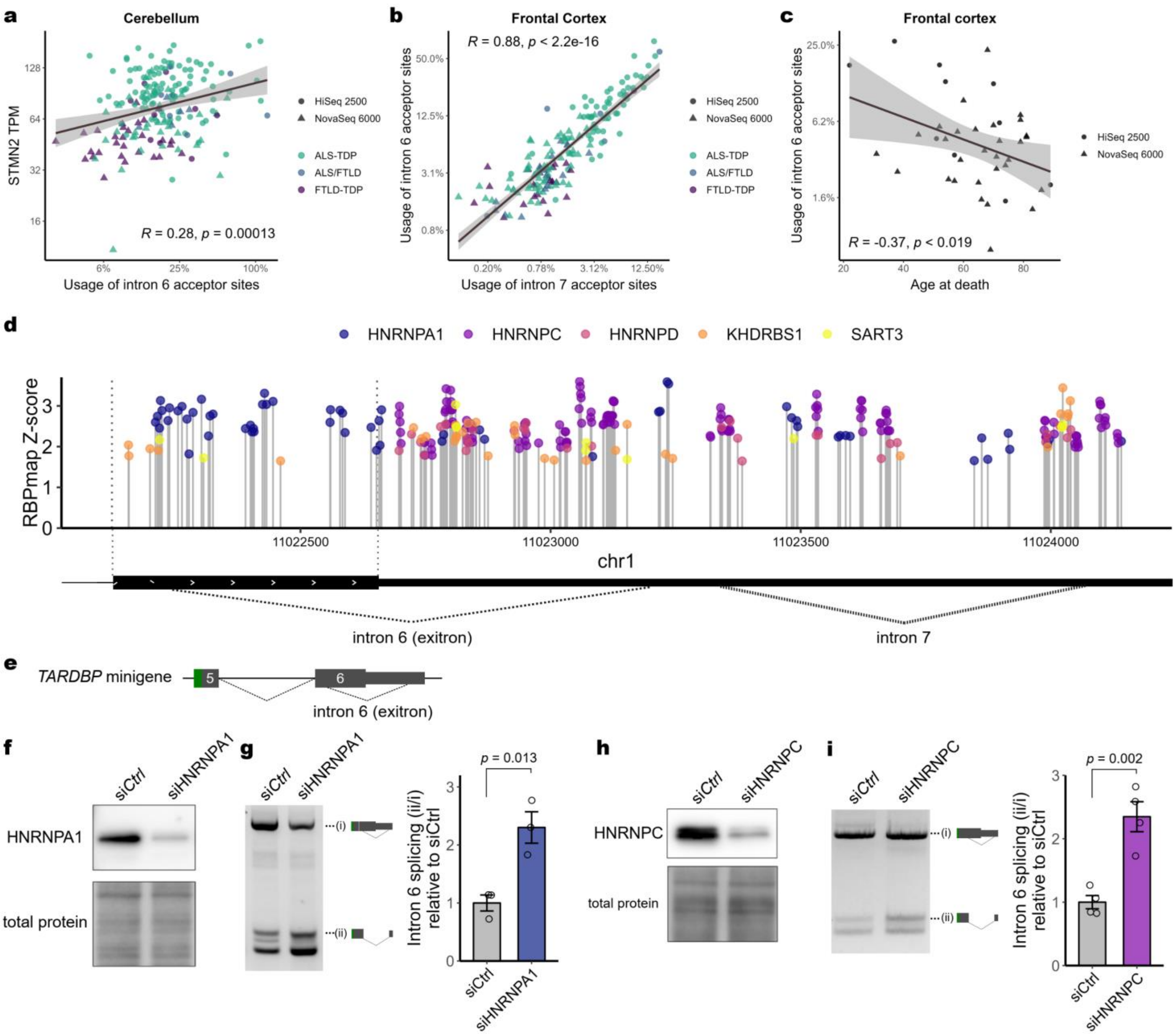


**Extended Data Fig. 1| Clinical relevance of *TARDBP* exitron splicing and identification of its regulatory RNA-binding proteins. a-c,** Further analysis of RNA-seq data from the NYGC ALS Consortium and frontal cortex of patients with TDP-43 proteinopathies (related to Fig. 1b-d). **a,** Association between *STMN2* expression and *TARDBP* exitron splicing in the cerebellum. **b,** Association between *TARDBP* intron 6 (exitron) and intron 7 splicing rates in the frontal cortex, demonstrating the coordinated regulation of these splicing events by TDP-43. **c,** Correlation between age at death and *TARDBP* intron 6 splicing rates in healthy controls. **d,** Position and Z-score of binding motifs in *TARDBP* for RBPs showing notable exitron splicing enhancement upon knockdown (Fig. 1e), as analyzed by RBPmap. **e,** Structure of the *TARDBP* minigene. **f-i,** Knockdown of HNRNPA1 (**f, g**) or HNRNPC (**h, i**) affects exitron splicing in the minigene. Data are mean ± SEM. Student's t-test.

#### Extended Data Fig.2

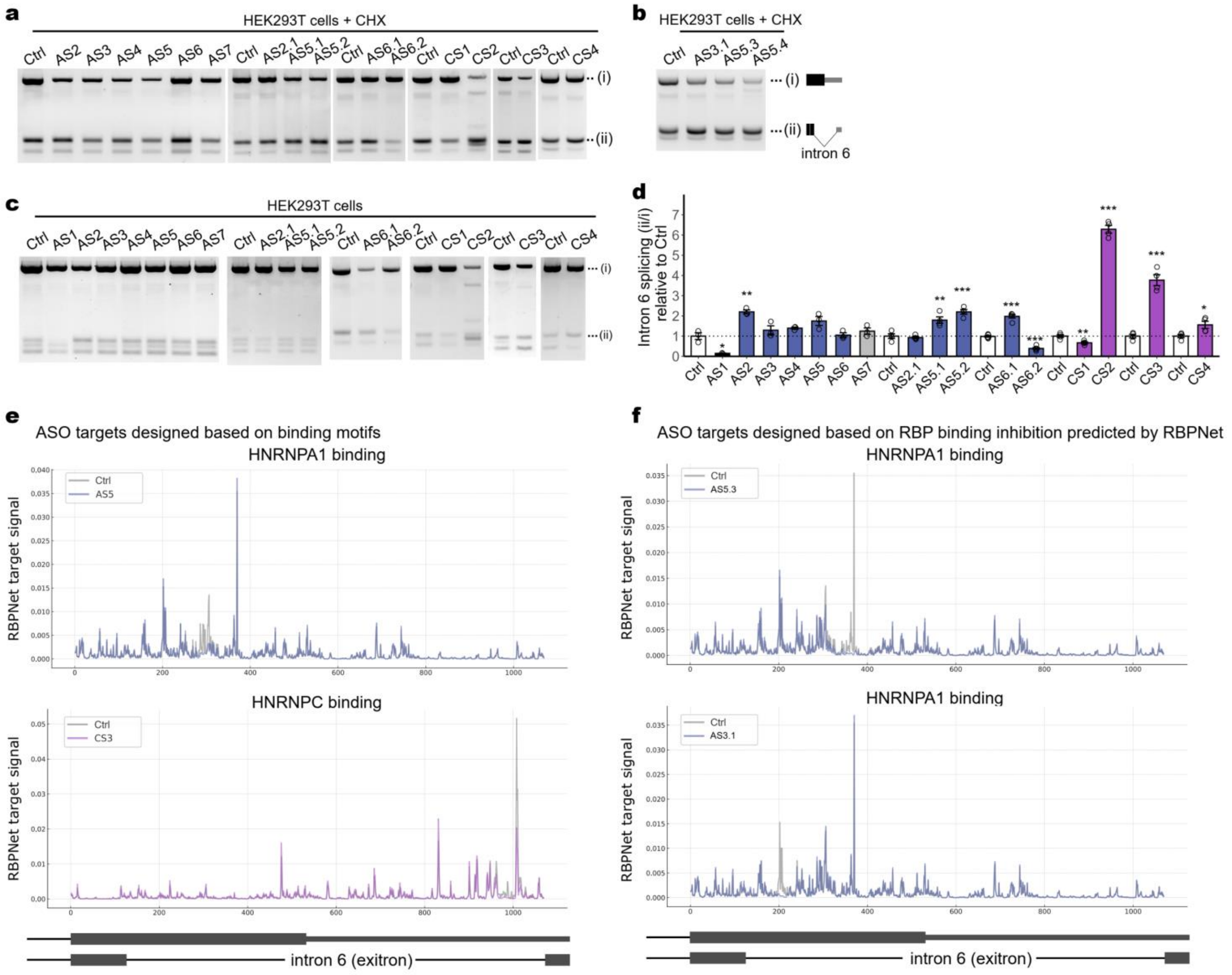


**Extended Data Fig. 2| ASOs targeting HNRNPA1 and HNRNPC binding promote *TARDBP* exitron splicing. a, b,** Intron 6 (exitron) splicing analysis in HEK293T cells transfected with ASOs and treated with cycloheximide (CHX) (related to Fig. 1h, i). **c, d,** Evaluation of intron 6 splicing in HEK293T cells transfected with ASOs without CHX treatment. AS1, an ASO that inhibits splicing, serves as an experimental control. AS1, targeting the exitron splicing donor site, served as an experimental control[11]. **e, f,** RBPNet predictions of HNRNPA1 and HNRNPC binding to the exitron with 'N' substitution, showing changes in binding index compared to control. Data are presented as mean ± SEM. Statistical analysis: Dunnett's test or Student's t-test vs control. *p < 0.05, **p < 0.01, ***p < 0.001.

#### Extended Data Fig.3

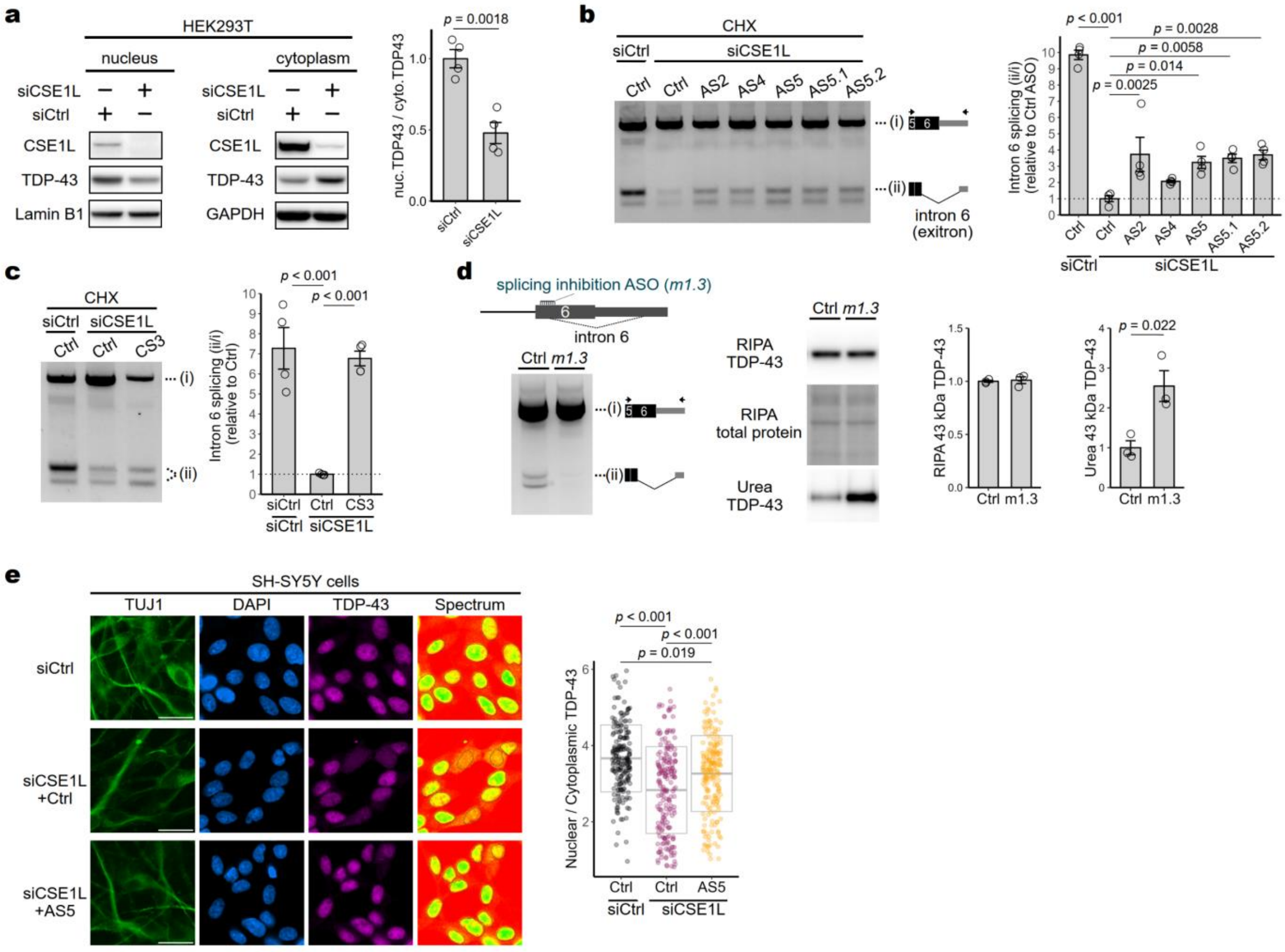


**Extended Data Fig. 3| Reduced *TARDBP* exitron splicing under impaired TDP-43 nuclear transport and ASO treatment effect. a,** Western blot of nuclear and cytoplasmic fractions from HEK293T cells with CSE1L/CAS knockdown. **b, c,** RT-PCR analysis of intron 6 splicing in HEK293T cells treated with siCSE1L or siCtrl and ASOs (control or specific targets) after CHX treatment. CSE1L knockdown reduces exitron splicing. ASOs targeting HNRNPA1 (**b**) and HNRNPC (**c**) binding regions restore splicing. Splicing rate shown as change relative to siCtrl with control ASO. **d,** Exitron splicing and TDP-43 levels in SH-SY5Y cells treated with splicing inhibition ASO (m1.3)[11]. **e,** Localization of TDP-43 in SH-SY5Y cells with CSE1L knockdown and AS5 treatment. Scale bar: 20 µm. Data are mean ± SEM. Student's t-test (**a,d**). Dunnett's test vs control ASO under siCSE1L (**b,c**). ANOVA followed by Tukey's multiple comparison test (**e**).

#### Extended Data Fig.4

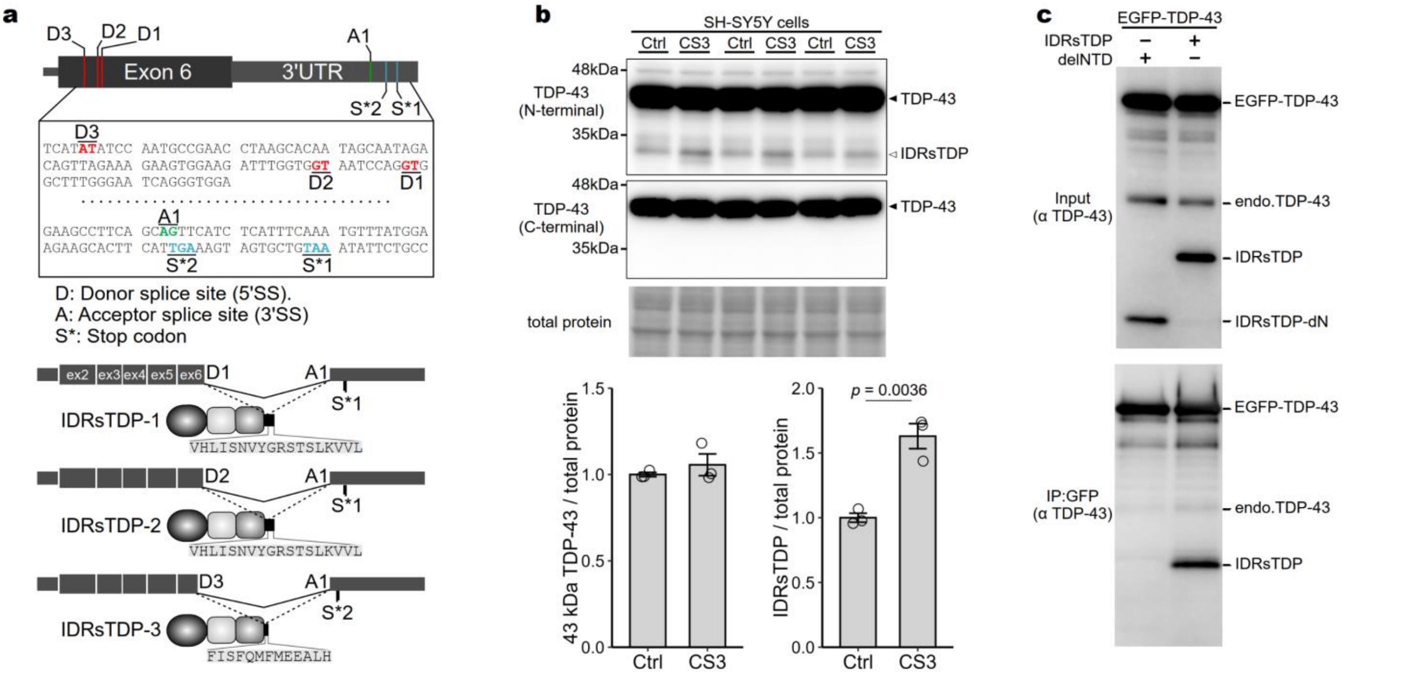


**Extended Data Fig.4| Exitron splicing is associated with expression of IDRsTDP, which binds TDP-43. a,** Schematic representation of *TARDBP* exon 6 and 3'UTR showing exitron splicing sites. D1-D3: Donor splice sites; A1: Acceptor splice site; S*1, S*2: Stop codons. Nucleotide sequences around splicing sites and amino acids added by splicing for IDRsTDP-1, IDRsTDP-2, and IDRsTDP-3 isoforms are shown. Representative splicing isoforms resulting from these splice sites are illustrated. **b**, Western blot of RIPA-soluble fractions from ASO-transfected SH-SY5Y cells using N-terminal TDP-43 antibody (10782-2-AP) and C-terminal TDP-43 antibody (TIP-TD-P09). CS3 increases IDRsTDP (~32 kDa band) detected by N-terminal antibody but not by C-terminal antibody (related to Figure 3a). Data are mean ± SEM. Student's t-test. **c,** Co-immunoprecipitation of EGFP-TDP-43 shows that the binding of IDRsTDP to TDP-43 is mediated by the NTD.

### Extended Data Fig.5

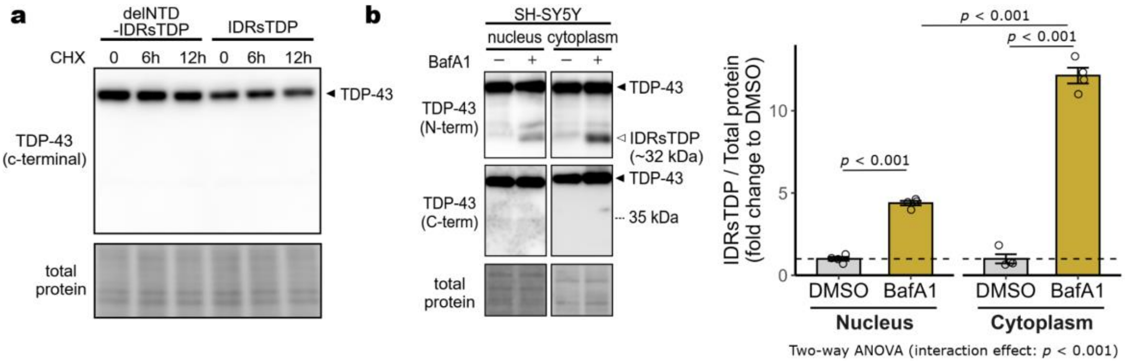


**Extended Data Fig.5| TDP-43 is degraded by autophagy via IDRsTDP. a,** Western blotting with a C-terminal TDP-43 antibody (TIP-TD-P09) does not detect IDRsTDP, IDRsTDP-dN, and the 25 kDa IDRsTDP fragment observed with an N-terminal TDP-43 antibody (related to Fig. 4a). **b,** Western blotting of nuclear and cytoplasmic fractions from SH-SY5Y cells treated with bafilomycin A1 shows an increase in IDRsTDP levels. Data are mean ± SEM.

### Extended Data Fig.6

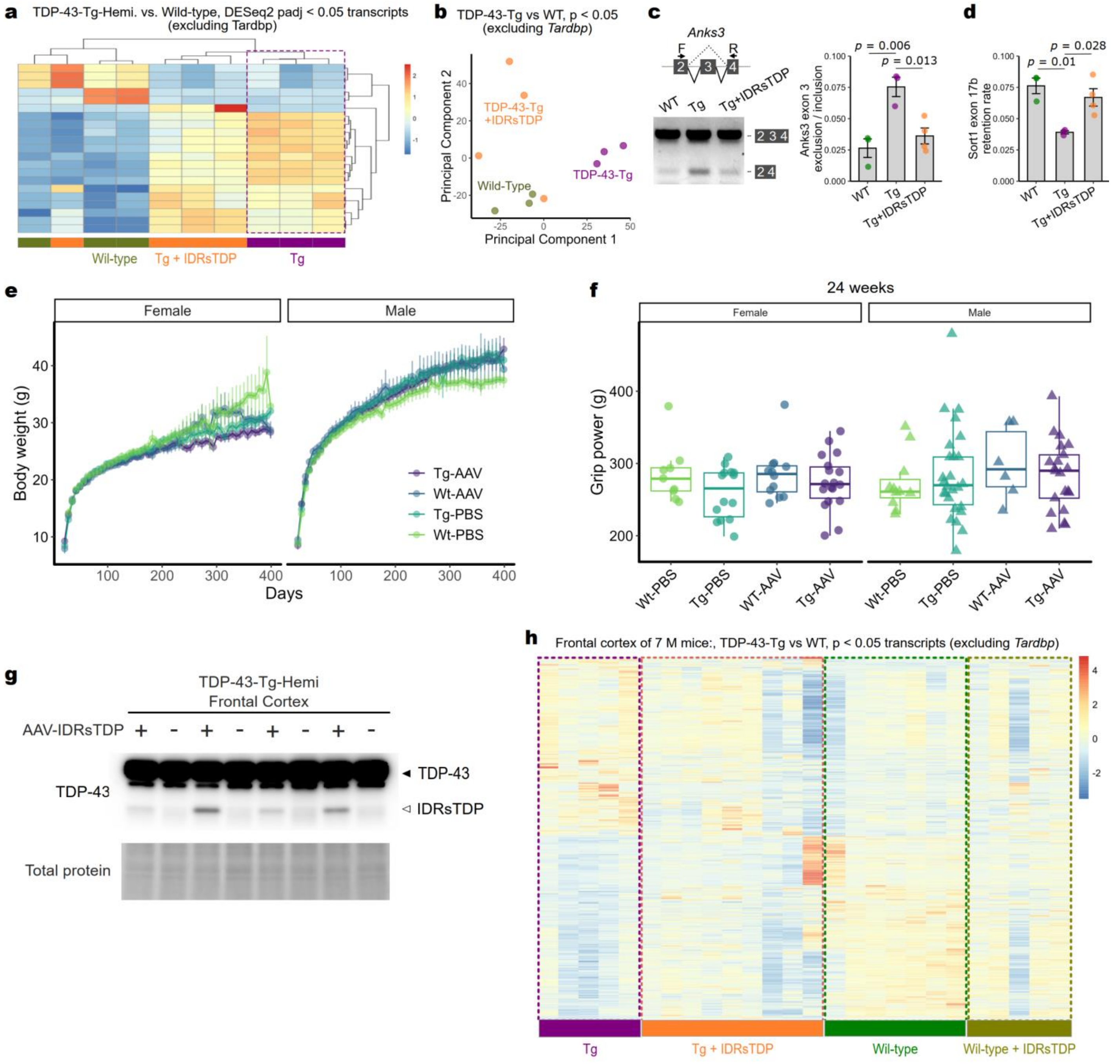


**Extended Data Fig.6| IDRsTDP suppresses RNA metabolic abnormalities caused by TDP-43 excess. a-d,** Analysis of TDP-43 Tg mice injected with AAV-sTDP (related to Fig 4f-i), including heatmap (**a**) and PCA (**b**) of transcripts, and evaluation of exon inclusion in Anks3 and Sort1 by RT-PCR (**c**) and ddPCR (**d**), respectively. ANOVA followed by Tukey's test (g, l, m). Mean ± SEM. **e-h,** Neonatal intracerebroventricular administration of AAV-IDRsTDP or PBS to WT or hemizygous TDP-43 Tg mice. **e,** Body weight changes monitored up to 400 days of age. **f,** Grip strength measured at 24 weeks of age (WT-PBS (Female) n = 9, Tg-PBS (Female) n = 14, WT-AAV (Female) n = 12, Tg-AAV (Female) n = 18, WT-PBS (Male) n = 11, Tg-PBS (Male) n = 26, WT-AAV (Male) n = 6, Tg-AAV (Male) n = 21). **g,** IDRsTDP expression in the frontal cortex at 7 months after injection of age. **h,** Heatmap showing transcripts with unadjusted p-values < 0.05 in the frontal cortex of TDP-43 Tg mice compared to WT controls at the 7-month time point.

#### Extended Data Fig.7

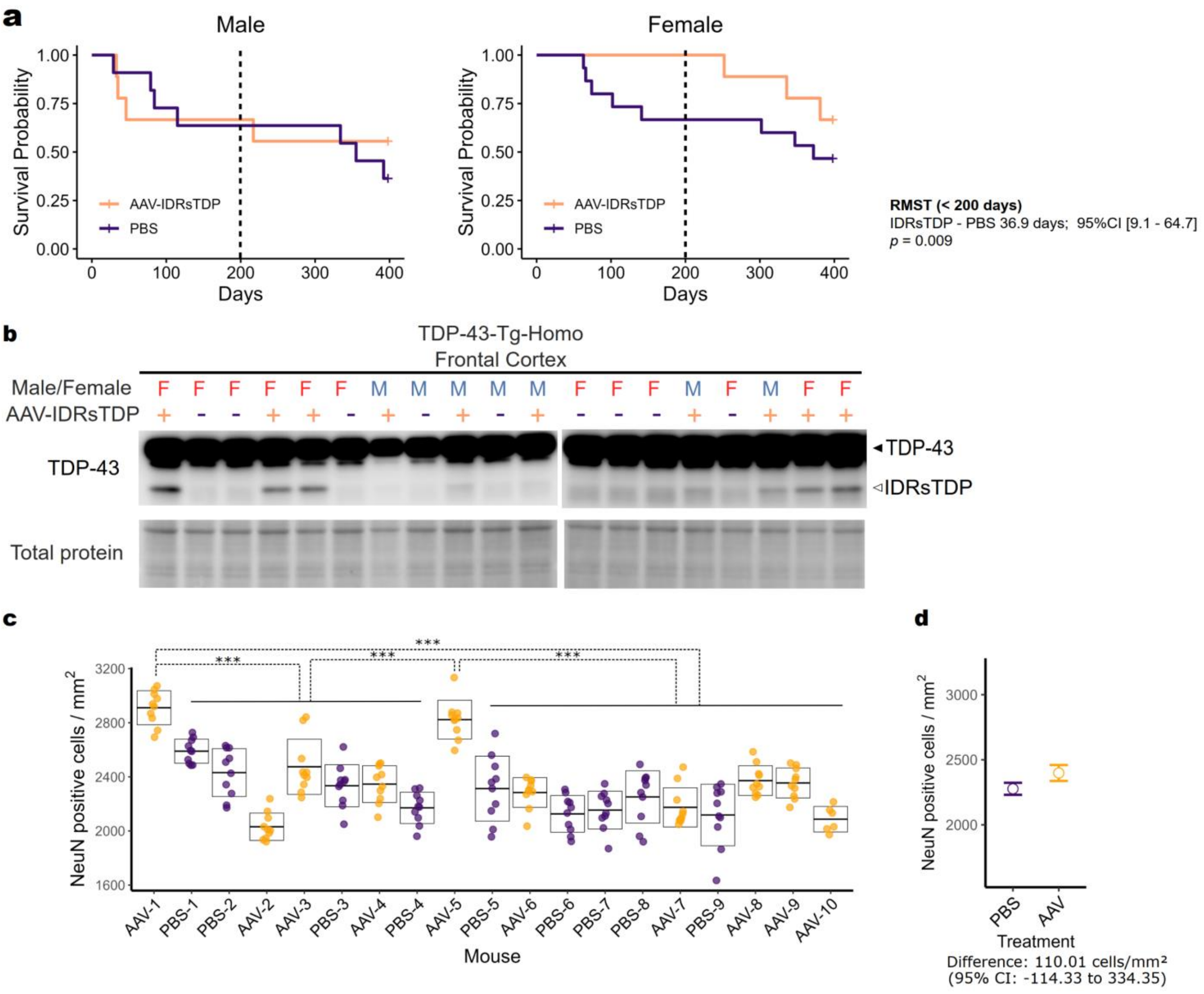


**Extended Data Fig.7| AAV-IDRsTDP improves survival in female TDP-43-Tg-Homo mice. a,** Survival curves for male (left) and female (right) TDP-43-Tg-homo mice treated with AAV-IDRsTDP or PBS. RMST analysis shows increased survival (<200 days) in AAV-IDRsTDP females ($p = 0.009$). **b,** Western blot of TDP-43 and IDRsTDP levels in the frontal cortex, normalized to total protein. **c,** Neuron (NeuN-positive cell) density in the frontal cortex of individual mice. Lines and boxes: mean ± SD. ANOVA with Tukey HSD (***$p < 0.001$). **d,** Means and 95% confidence Intervals for each treatment group. linear mixed-effects model analysis. The model accounted for within-mouse variability as a random effect.

### Extended Data Fig.8

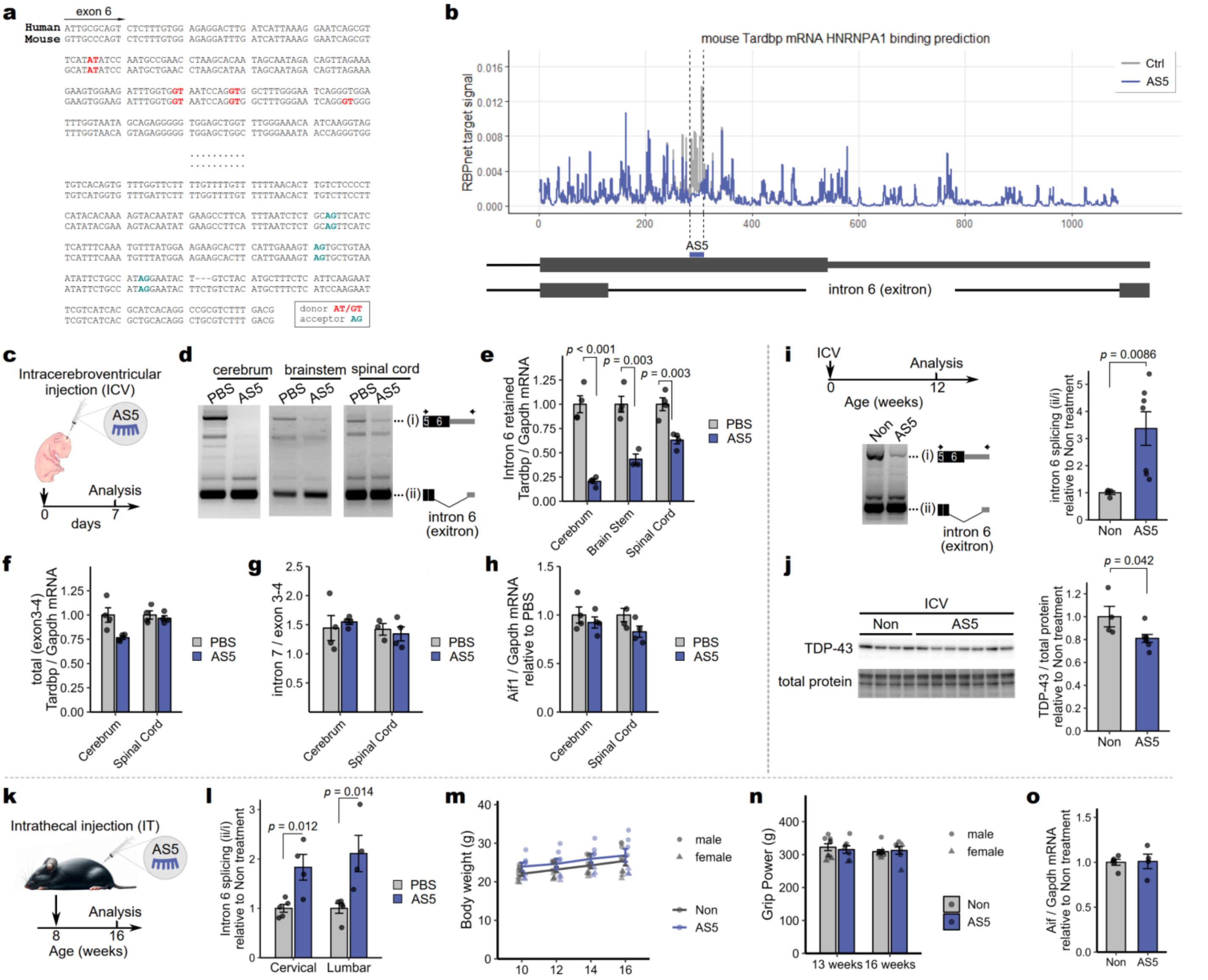


**Extended Data Fig.8| AS5 promotes exitron splicing in wild-type mice. a,** Sequences containing exon 6 to exitron of the *TARDBP* gene. The upper row shows human and the lower row shows mouse sequences. Red indicates donor sites and light green indicates acceptor sites of the exitron. **b,** Prediction of HNRNPA1 binding to mouse *Tardbp* exon 6 to exitron by RBPNet (gray line). The blue line shows the predicted signal when the AS5 target sequence is substituted with N to mimic inhibition of HNRNPA1 binding. **c-h,** Intracerebroventricular injection (ICV) of 4mM AS5 (2 µl) in neonatal mice, with RNA analysis of cortex, brainstem, and spinal cord tissues after 1 week. **d,** AS5 promoted intron 6 (exitron) splicing. **e,** Real-time PCR of mRNAs retaining intron 6. **f,** Quantification of total *TARDBP* mRNA using exon 3-4 primers. **g,** Comparison of intron 7-containing RNA to total *TARDBP*. **h,** Aif mRNA expression by qPCR. **i,j,** ICV of 4mM AS5 (3 µl) in neonatal mice, with analysis performed 12 weeks post-injection. **i,** Analysis of intron 6 splicing. **j**, TDP-43 protein levels. **k-o,** Intrathecal injection of 4mM AS5 (15 µl) AS5 in 8-week-old wild-type mice, with body weight, grip strength, and spinal cord analysis after 8 weeks. **l,** RT-PCR showing enhanced intron 6 splicing (related to Fig. 5c). **m,** Body weight measured every 2 weeks. **n,** Grip strength at 13 and 16 weeks. **o,** Aif mRNA expression by qPCR. Data are mean ± SEM.

### Extended Data Fig.9

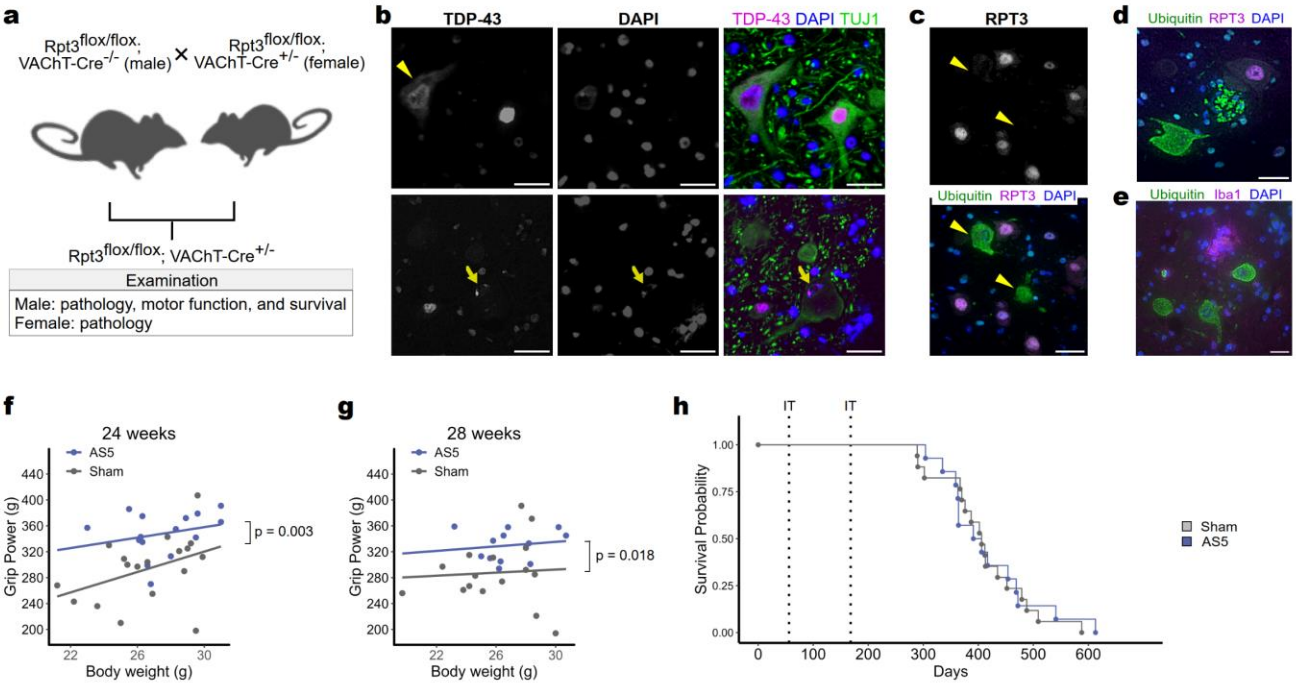


**Extended Data Fig.9| Analysis of *Rpt3*$^{\text{flox/flox}}$;*VAChT-Cre*$^{+/-}$ mice treated with ASO after the appearance of pathology.** **a,** Reproduction manner of *Rpt3*$^{\text{flox/flox}}$;*VAChT-Cre*$^{+/-}$ mice. **b,** Representative image illustrating mislocalization of TDP-43 (upper panel) and reduction and aggregation of nuclear TDP-43 (lower panel). **c,** Representative image illustrating ubiquitin positivity in RPT3-negative spinal motoneurons of 8-week-old *Rpt3*$^{\text{flox/flox}}$;*VAChT-Cre*$^{+/-}$ mice. **d,** Representative image displaying traces of death in ubiquitin-positive motoneurons. **e,** Representative image depicting traces of death in ubiquitin-positive motor neurons, surrounded by Iba1-positive microglia. **f,** Relationship between body weight and grip strength at 24 and 28 weeks in mice that received intrathecal (IT) injections of AS5 or sham surgery at 8 and 24 weeks after the onset of pathology. Grip strength is significantly higher in the AS5-treated group, independent of body weight. Statistical significance was determined by ANCOVA. **g,** Final survival duration. Scale bar: 30 μm (**c-e**) or 40 μm (**b**).

**Supplementary Table 1| Primer and probe sequence.**

| Name | Sequence (5' to 3') | Experimental Purpose |
|---|---|---|
| *TARDBP*_Exitron_F | GCGCTGTACAGAGGACATGAC | Amplification of *TARDBP* exitron region |
| *TARDBP*_Exitron_R | GCCTGTGATGCGTGATGA | Amplification of *TARDBP* exitron region |
| *TARDBP*_Minigene_F | AAGACGGCATACGAGAC | *TARDBP* minigene exitron splicing |
| *TARDBP*_Minigene_R | GCCTGTGATGCGTGATGA | *TARDBP* minigene exitron splicing |
| *STMN2*_F | AGCTGTCCATGCTGTCACTG | *STMN2* expression quantification |
| *STMN2*_R | GTGGCTTCAAGATCAGCTC | *STMN2* expression quantification |
| *TARDBP*_Splice_V1_F | AAAGAAGTGGAAGATTTGGTGTTC | *TARDBP* splice variant 1 |
| *TARDBP*_Splice_V2_F | GAAGATTTGGTGGTAATCCAGTTC | *TARDBP* splice variant 2 |
| *TARDBP*_Splice_V1V2_R | GCCTGTGATGCGTGATGA | *TARDBP* splice variants 1 and 2 |
| *TARDBP*_Unspliced_F | TGTCACAGTGTTTGGTTCTTTTG | Unspliced *TARDBP* isoform |
| *TARDBP*_Unspliced_R | AGCGGATAAAAAATGGGACAC | Unspliced *TARDBP* isoform |
| Mouse_*Tardbp*_Ex1_F | GCCGTTCTGTCCTTCATCTG | Mouse *Tardbp* |
| Mouse_*Tardbp*_Ex1_R | CAGGCTCTTCGGCGTATAGA | Mouse *Tardbp* |
| Human_TDP43_Ex5-6_F | CTGCGGGAGTTCTTCTCTCA | Human TDP-43Tg genotyping |
| Human_TDP43_Ex5-6_R | CGCAATCTGATCATCTGCAA | Human TDP-43Tg genotyping |
| Mouse_*Tardbp*_Ex1_Probe | /56-FAM/CAGTTTTTC/ZEN/AGACCCAGCTGTTTTCAT/3IABkFQ/ | Mouse *Tardbp* |
| Human_TDP43_Ex5_Probe | /5HEX/CCATTCAGG/ZEN/GCCTTTGCCTTTGTT/3IABkFQ/ | Human *TARDBP* genotyping |
| Myc_tag_F | AGTAGTGCTGGAACAAAAACTCATC | Vector gene detection |
| WPRE_R | TCCAGAGGTTGATTATCGGAATTC | Vector gene detection |
| Mouse_*Tardbp*_Exitron_F | CAGAGCTTTTGCCTTCGTCA | Mouse *Tardbp* exitron splicing |
| Mouse_*Tardbp*_Exitron_R | CAAAGACGCAGCCTGTGC | Mouse *Tardbp* exitron splicing |
| Mouse_*Tardbp*_Retained_F | AGGTGGCTTTGGGAATCAG | Mouse *Tardbp* exitron-retained mRNA |
| Mouse_*Tardbp*_Retained_R | CACCAAAGTTCATCCCTCCA | Mouse *Tardbp* exitron-retained mRNA |
| Mouse_*Tardbp*_Total_F | AACTGAGCAGGATCTGAAAGAC | Total mouse *Tardbp* mRNA |
| Mouse_*Tardbp*_Total_R | CGAACAAAGCCAAACCCTTTC | Total mouse *Tardbp* mRNA |
| Mouse_*Tardbp*_Total_Probe | 56-FAM/TGGAGAGGT/ZEN/TCTTATGGTTCAGGTCA/3IABkFQ | Total mouse *Tardbp* mRNA |
| Mouse_*Tardbp*_Intron7_F | TGCTGTATGGTGTGTGTTCTC | Mouse *Tardbp* intron 7-retained mRNA |
| Mouse_*Tardbp*_Intron7_R | CCACAAGCTCAGTCCATGTT | Mouse *Tardbp* intron 7-retained mRNA |
| Mouse_*Tardbp*_Intron7_Probe | 56-FAM/AGTGTGGGA/ZEN/ACGTGAACTGAAGCT/3IABkFQ | Mouse *Tardbp* intron 7-retained mRNA |
| *Aif*_F | ACGAACCCTCTGATGTGGTC | *Aif* expression quantification |
| *Aif*_R | CGGGATGGAAGAGAGAGGA | *Aif* expression quantification |
| *Gapdh*_F | TGTGTCCGTCGTGGATCTGA | Internal control |
| *Gapdh*_R | TTGCTGTTGAAGTCGCAGGAG | Internal control |
